# Target-specific control of olfactory bulb periglomerular cells by GABAergic and cholinergic basal forebrain inputs

**DOI:** 10.1101/2021.07.06.451255

**Authors:** Didier De Saint Jan

**Affiliations:** Institut des Neurosciences Cellulaires et Intégratives, Centre National de la Recherche Scientifique, Unité Propre de Recherche 3212, Université de Strasbourg, 67084 Strasbourg, France

**Keywords:** acetylcholine, GABA, olfactory bulb, basal forebrain, periglomerular cells

## Abstract

The olfactory bulb (OB), the first relay for odor processing, receives dense GABAergic and cholinergic long-range projections from basal forebrain (BF) nuclei that provide information about the internal state and behavioral context of the animal. However, the targets, impact and dynamics of these afferents are still unclear. I studied how BF synaptic inputs modulate activity in diverse subtypes of periglomerular (PG) interneurons using optogenetic stimulation and loose cell-attached or whole-cell patch-clamp recording in OB slices from adult mice. GABAergic BF inputs potently blocked PG cells firing except in a minority of calretinin-expressing cells in which GABA release elicited spiking. Parallel cholinergic projections excited a previously overlooked PG cell subtype via synaptic activation of M1 muscarinic receptors. Low frequency stimulation of the cholinergic axons drove persistent firing in these PG cells thereby increasing tonic inhibition in principal neurons. Taken together, these findings suggest that modality-specific BF inputs can orchestrate inhibition in OB glomeruli using multiple, potentially independent, inhibitory or excitatory target-specific pathways.

## Introduction

Basal forebrain (BF) nuclei innervate many regions of the brain including the entire neocortex, hippocampus, amygdala, thalamus and hypothalamus with diffuse long-range projections releasing GABA, ACh and, more rarely, glutamate. These projections are thought to provide cues about the behavioral context and internal state of the animal. They modulate multiple synaptic, cellular and network processes at a variety of temporal and spatial scales thereby regulating sensory perception, metabolic functions such as food intake, brain states and important cognitive functions including attention, arousal, memory or learning (Picciotto et al. 2012, Ballinger et al. 2016).

The olfactory bulb (OB), the first region that processes olfactory information in the brain, receives massive cholinergic and GABAergic BF projections that principally originate in the nucleus of the horizontal limb of the diagonal band of Broca (HDB) and in the magnocellular preoptic nucleus (MCPO) (Zaborszky et al. 1986). Cholinergic signaling within the OB modulates olfactory learning and memory (Ravel et al. 1994, Devore et al. 2012, Devore et al. 2014, Ross et al. 2019), odor discrimination (Doty et al. 1999, Mandairon et al. 2006, Chaudhury et al. 2009, Li and Cleland 2013, Smith et al. 2015, Chan et al. 2017), odor habituation (Ogg et al. 2018) or social interactions (Suyama et al. 2021). How ACh modulates each of these behavioral demands is less clear. Cholinergic axons innervate all layers of the OB and preferentially form synapses on inhibitory interneurons (Kasa et al. 1995, Hamamoto et al. 2017). Although several classes of neurons, including principal neurons, express diverse nicotinic and muscarinic receptors (Castillo et al. 1999, Liu et al. 2015, Smith et al. 2015. Reviewed in Brunert and Rothermel 2021), deep short axon cells are the only known synaptic targets of BF cholinergic neurons to date (Case et al. 2017). Cholinergic neurons innervating deep short axon cells release both ACh and GABA. Co-transmission of ACh and GABA at BF axon terminals is systematic in the hippocampus (Takacs et al. 2018) but seems target-specific in the cortex (Saunders et al. 2015, Desikan et al. 2018). Thus, what class of OB neurons is modulated by BF cholinergic projections during a specific olfactory task, what type of ACh receptors are recruited, how endogenous ACh activates these receptors (volume or synaptic transmission?) and is GABA always co-transmitted with ACh in the OB are still unresolved questions.

BF GABAergic afferents innervate all layers of the OB at least as densely as cholinergic axons but only few studies have examined their function in odor processing (Nunez-Parra et al. 2013, Bohm et al. 2020). This is a difficult question because GABAergic projections to the OB likely arise from several populations of BF neurons (Sanz Diez et al. 2019). Moreover, electrophysiological evidence indicate that BF GABAergic afferents innervate several types of GABAergic interneurons. These targets include granule cells and PG cells which inhibit principal neurons (Nunez-Parra et al. 2013, Sanz Diez et al. 2019, Hanson et al. 2020, Villar et al. 2021) and deep short-axon cells which inhibit granule cells and PG cells (Case et al. 2017, Sanz Diez et al. 2019). Thus, depending on their target BF GABAergic inputs may inhibit or disinhibit principal neurons. Finally, the effect on principal neurons may be hard to predict as GABA is excitatory in some PG cells (Parsa et al. 2015).

In such a complex context, an important step towards clarifying the influence and function of the BF inputs in the OB is to investigate the connections, temporal dynamics and impact of each BF pathway. BF axons are particularly abundant in the glomerular layer of the bulb where they innervate three classes of PG cells with target-specific release properties and IPSC time course (Sanz Diez et al. 2019). In addition, muscarinic receptors exert a strong control on glomerular inhibition (Liu et al. 2015) suggesting that PG cells are a likely target of cholinergic axons. Here, I examined how synaptic release of GABA and ACh from BF fibers modulates the activity of the various PG cell subtypes. The results demonstrate that PG cell subtypes are differently controlled by BF afferents and reveal that a previously overlooked PG cell subtype is a central player in mediating BF muscarinic modulation of glomerular inhibition.

## RESULTS

### Optogenetic stimulation of BF axons induces heterogeneous responses in PG cells

ChR2 fused with eYFP was targeted to BF neurons by injecting a viral construct into the HDB/MCPO of dlx5/6-Cre mice (Fig.1A). As reported in our previous study (Sanz Diez et al. 2019), this induced the expression of ChR2-eYFP in several types of GABAergic neurons as well as in cholinergic neurons in the BF. Axonal projections of these neurons also expressed ChR2-eYFP and densely innervated the OB (Fig.1B).

**Figure 1:**
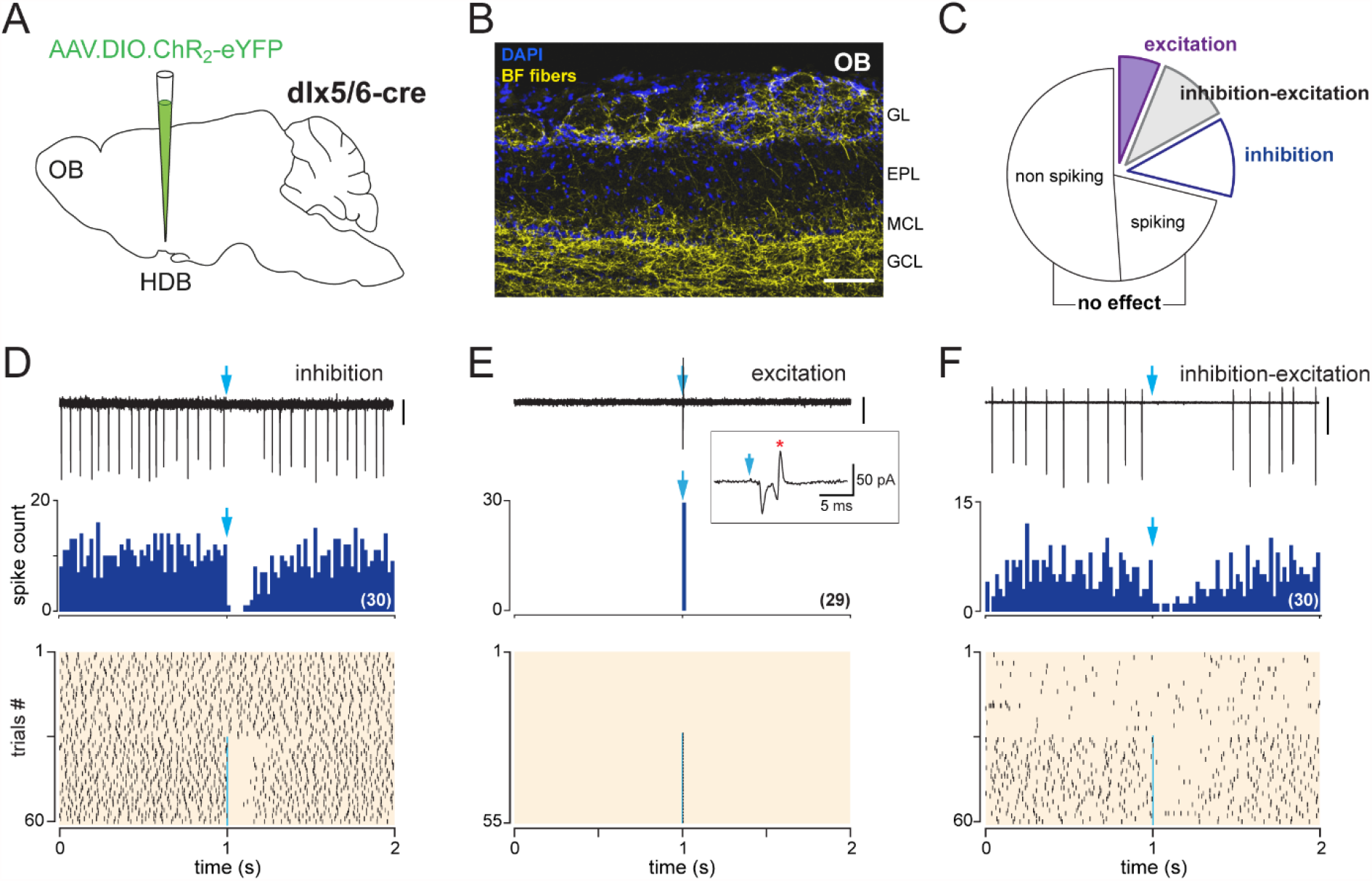
BF inputs have various impacts on PG cells activity. **A**, Schematic of the virus injection in the HDB/MCPO of dlx5/6-cre mice. **B**, ChR2-eYFP expressing BF axons (yellow) in a sagittal section of the OB. DAPI staining (blue) delimits the different layers (GL: glomerular layer; EPL: external plexiform layer; MCL: mitral cell layer; GCL: granule cell layer). Scale bar 100 µm. **C**, Probability of occurrence of the various types of LCA responses induced on PG cells firing by a single optogenetic stimulation of the BF axons. **D**, Example of an inhibitory response. In this cell, photostimulation of the BF fibers (blue arrow, 1 ms flash, 490 nm) transiently inhibited spiking. Top, a representative 2 s-long LCA recording episode (scale bar 20 pA). Middle, cumulative PSTH (bin size 20 ms) in the same cell for 30 consecutive episodes with stimulation. Bottom, raster plot showing spiking activity in control condition (episodes 0-30) and while BF axons were photostimulated once per episode with a single flash (episodes 31-60, blue line). Each tick indicates an action potential. **E**, example of an excitatory response where BF fibers photostimulation (episode 27-55) induced an inward current and a spike (red star in the inset). Scale bar 30 pA. Corresponding PSTH and raster plot, same as in D. **F**, example of a dual inhibition-excitation response where BF axons photostimulation inhibited spiking after the flash but also induced an increase in baseline firing rate (see raster plot). Scale bar 200 pA. PSTH and raster plot, same as in D and E.

Next, I used loose cell-attached (LCA) recording in acute OB slices from these mice (thereafter called dlx5/6 mice) to monitor spiking activity in randomly chosen PG cells while BF axons were periodically photostimulated every 2 seconds (0.5 Hz) using a single brief (1 ms) flash of blue light. Post stimulus spiking activity was affected in 29% of the cell tested (109/383, Fig.1C). This fraction is small compared to the proportion of PG cells that receive direct BF GABAergic inputs (around 80%)(Sanz Diez et al. 2019). However, more than half of the cell tested were silent or fired only rarely (i.e. <1 Hz) and the impact of a BF input may simply be unnoticeable in these cells using LCA spike recording. Calretinin (CR)-expressing PG cells, the most abundant PG cells, cannot fire or fire at low rate (Fogli Iseppe et al. 2016, Benito et al. 2018) and thus fall into this category although they do receive large GABAergic BF inputs (Sanz Diez et al. 2019). In contrast, spontaneously spiking cells that did not respond to the stimulation (20% of the cells, n=78) are likely PG cells that do not receive BF synaptic input such as olfactory sensory neurons (OSN)-innervated type 1 PG cells (Sanz Diez et al. 2019).

In our previous report, we found that BF axons mediate target-specific GABAergic IPSCs onto three subtypes of PG cells (Sanz Diez et al. 2019). These targets are all type 2 PG cells, i.e. they are not directly connected to OSN. Thus, not surprisingly, three clearly distinct types of response were evoked by the photostimulation of BF axons. However, the nature of these responses was unexpected (Fig.1D-F). 12% of the cells tested (45/383) responded with a transient inhibition of spiking immediately after the stimulation (Fig.1D), as expected for a classical inhibitory input. In contrast, in 6% of cells, photostimulation of the BF input elicited spiking (22/383 cells, Fig.1E). Finally, a dual inhibition-excitation response was evoked in 11% of cells (42/383 cells, Fig.1F). In these cells, photostimulation of BF axons transiently inhibited spiking and in parallel increased baseline firing rate. The principal focus of this study is on this unexpected dual response (Fig.2-8), although the two others kind of responses are also examined in Fig.8 and Fig.9.

**Figure 2:**
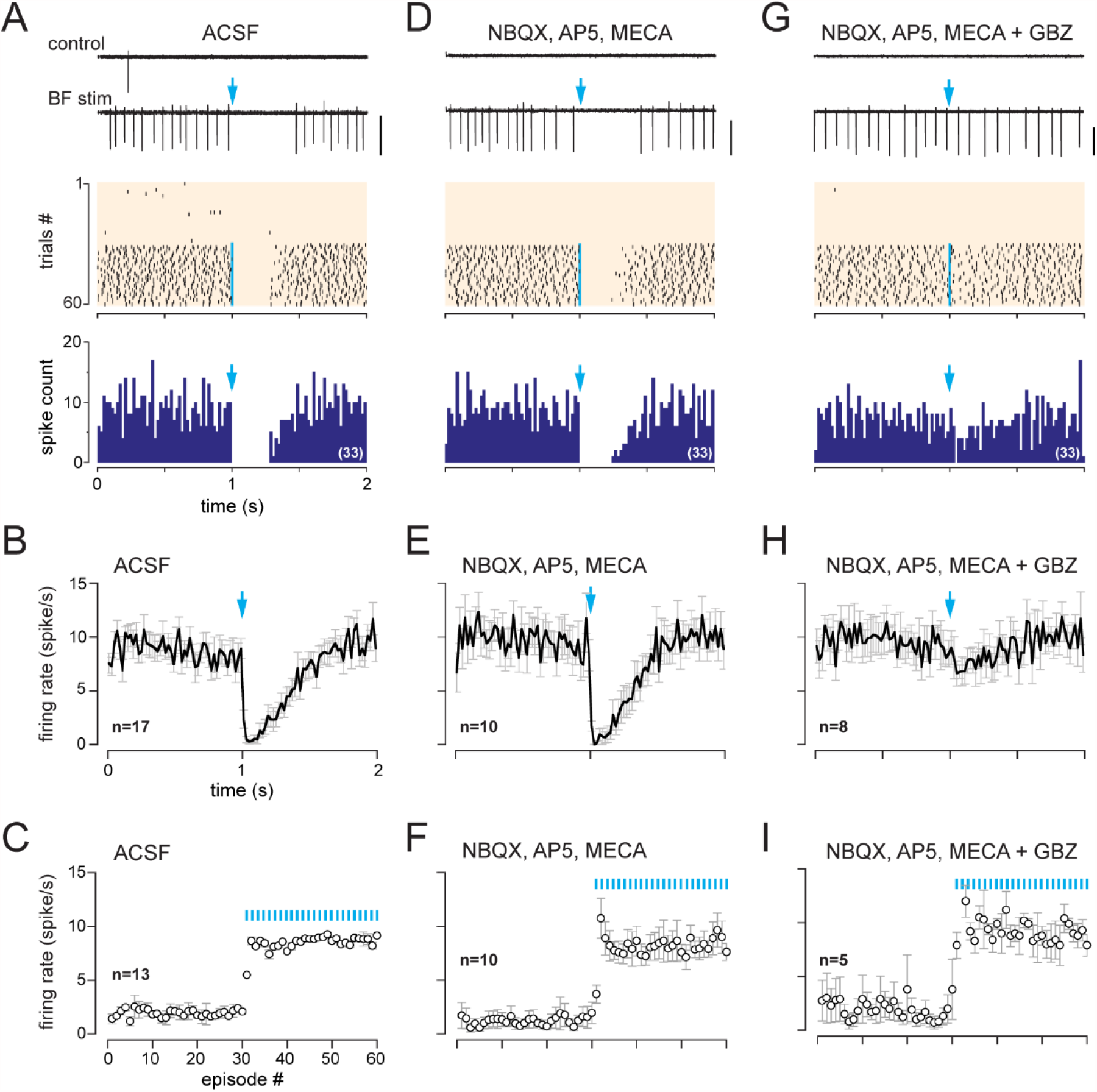
A monosysnaptic GABAergic input mediates the inhibitory component of the dual inhibition-excitation response. **A**, Two representative traces (scale bar 100 pA) showing spiking activity in control condition and while BF axons were photostimulated with a single flash per episode (blue arrow). Middle, corresponding raster plot. Photostimulation started at episode 31 (blue line). Bottom, cumulative PSTH (bin size 20 ms) in the same cell for the trials with photostimulation (blue arrow). **B**, average firing rate per bin (20 ms) and per episode for 17 cells (+/- sem, grey bars) while BF axons were photostimulated with a single flash (at blue arrow). **C**, average firing frequency per episode (2 s each) for 13 cells recorded in ACSF in control condition (no light, episodes 1-30) and during photostimulation of the BF afferents once per episode (31-60). **D-F**, Same as in A-C in the presence of NBQX (10 µM), D-AP5 (50 µM), and mecamylamine (MECA, 50 µM). Traces, raster plot and PSTH in D are from the same cell as in A. **G-I**, Same as in A-C when gabazine (GBZ, 5 µM) was added to the cocktail of blockers. Traces, raster plot and PSTH in G are from the same cell as in A and D.

### GABA and ACh release from BF axons mediate dual inhibition-excitation responses

Dual inhibition-excitation responses were found in PG cells with a low spiking activity in control condition (range 0-4 Hz, mean 1.95 ± 1.6 Hz, n=13). As illustrated in Fig.2A, baseline firing rate sharply increased as soon as 0.5 Hz photostimulations of the BF fibers started. Cells that were silent started to fire regularly whereas cells that had a low and irregular firing activity adopted a higher and more regular spiking rate (mean 8.6 ± 3 Hz, n=13, Fig.2C). This sustained spiking regime was maintained throughout the trials with stimulation (>1 min)(Fig.2A, 2C). In addition, spiking was transiently blocked immediately after the flash for 391 ± 142 ms (n=21, Fig.2A, 2B). Photo-evoked spiking inhibition and increase in baseline firing rate both persisted in the presence of NBQX (10 µM), D-AP5 (50 µM) and mecamylamine (50 µM)(n=10, Fig.2D-F). These antagonists inhibit AMPA, NMDA and nicotinic ACh receptors, respectively. As these compounds block any excitation of intermediate neurons that would modulate PG cells, both spiking inhibition and increase in baseline spiking appear to be caused by a direct monosynaptic pathway. In contrast, the GABA_A_ receptor antagonist gabazine (5 µM) blocked spiking inhibition but did not prevent the increase of basal firing (n=8, Fig.2G-I). This suggests that a direct BF GABAergic IPSC mediates spiking inhibition whereas the excitatory component is caused by a different pathway.

Next, I used a longer (15-20 s) interval between each flash to examine the time-course of the excitatory component of the dual inhibition-excitation responses. This protocol revealed that a single photostimulation elicits a reliable and long-lasting increase in firing rate (Fig.3A, 3B). Spike rate increased almost 4 fold compared to baseline activity (from 2.9 ± 2.7 Hz before the flash to 10.9 ± 4.6 Hz 1 s after the flash, n=32, Fig.3B, 3D), peaked 1-2 s after the flash and returned to baseline frequency after about 10 s (Fig.3B). This long-lasting photo-evoked excitation was stable for multiple stimulations and always followed the gabazine-sensitive phase of spike inhibition in all but one cell (Fig.3A, 3B. Fig.3-figure supplement 1). Evoked excitation persisted in the presence of NBQX, D-AP5 and mecamylamine (spike frequency increased from 1.9 ± 1.7 Hz before the flash to 9.4 ± 3.0 Hz after the flash, n=14, Fig.3D) and was not affected by gabazine (n=3, not shown). In contrast, photo-evoked excitation was totally blocked by atropine (10 µM, n=7) or scopolamine (10 µM, n=2), two non-selective antagonists of metabotropic muscarinic ACh receptors (mAChRs)(Fig.3C, 3D). Thus, synaptic release of ACh from BF axon terminals strongly excites a subset of PG cells via the activation of mAChRs. Baseline firing frequency was also strongly reduced or totally abolished in the presence of a mAChR antagonist in 7/9 cells, making it difficult to evaluate if photo-evoked inhibition of spiking was affected or not. However, in the two cells that still spontaneously fired in the presence of atropine or scopolamine, the inhibitory component seemed to persist (not shown). All together, these results indicate that a subset of PG cells receive both ACh and GABA inputs from BF axons. GABA release inhibits spiking through GABA_A_ receptors whereas ACh release activates mAChRs and produces a previously unreported long-lasting excitation.

**Figure 3:**
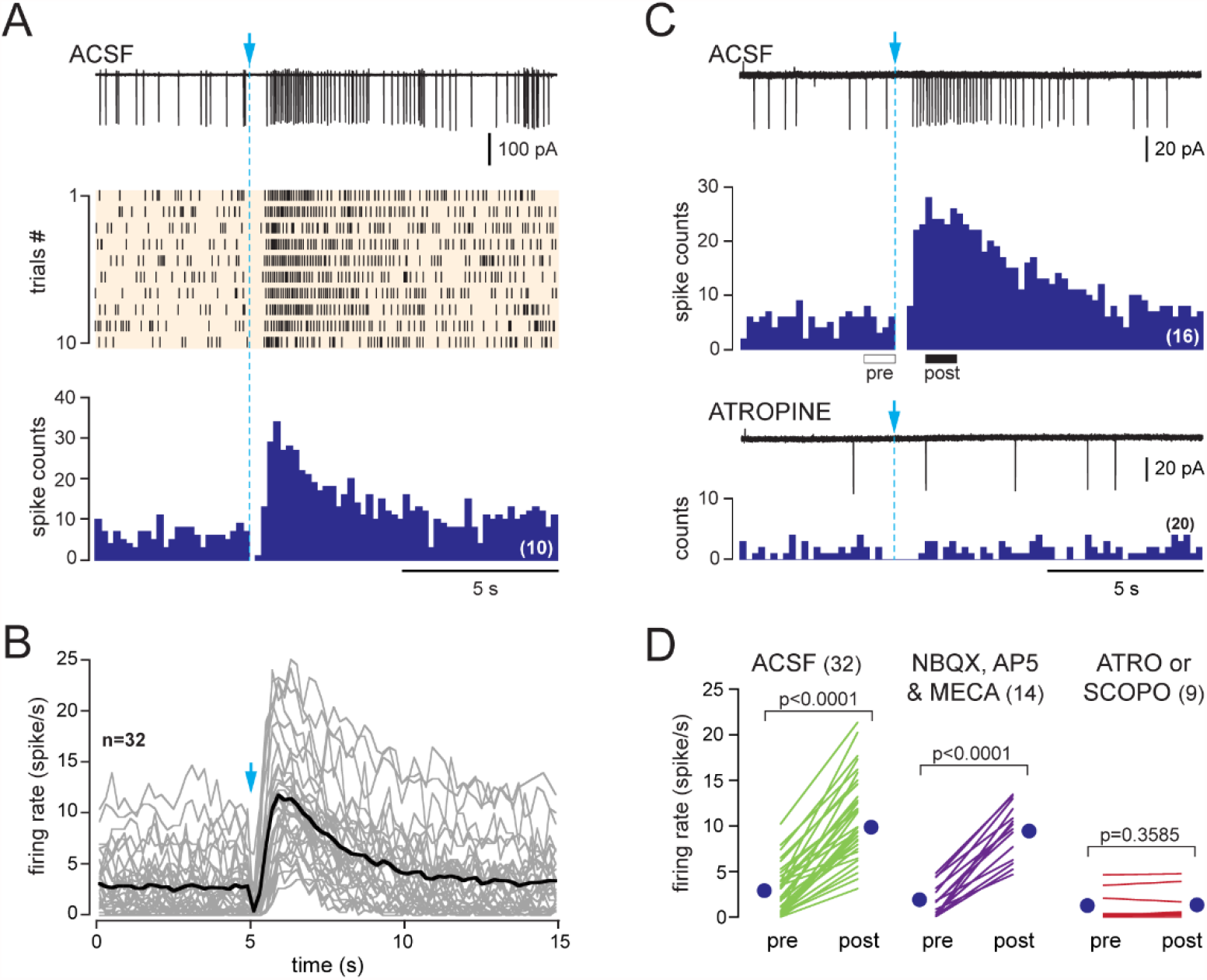
A single BF axons stimulation evokes a long-lasting muscarinic excitation in PG cells with dual inhibition-excitation responses. **A**, Representative spiking response, raster plot and cumulative PSTH (10 consecutive sweeps, 200 ms/bin) of a typical dual inhibition-excitation response evoked by a single photostimulation of BF fibers and recorded over 15 s in a PG cell from a dlx5/6 mouse. Only one cell responded with a long-lasting excitation that was not preceded by an inhibitory component (Fig.3-figure supplement 1). **B**, Average spiking frequency per bin (200 ms) and per episode. Each grey line corresponds to a cell. The black line is the ensemble average. Photostimulation at blue arrow. **C**, The non-selective mAChR antagonist atropine (10 µM) blocked BF-evoked excitation. Pre and post boxes below the control PSTH indicate the time periods that were compared in D. **D**, Firing rate before (pre) and after (post) photostimulation of BF axons in ACSF (green), in the presence of NBQX, D-AP5 and mecamylamine (violet) or in the presence of the muscarinic receptor antagonist atropine (n=7) or scopolamine (n=2)(red). Each line indicates a cell. Blue circles indicate means. Paired t-test.

### GABA and ACh are released by separate BF axons

Although observed in only one case, the example of a BF-evoked muscarinic excitation lacking the GABAergic component (Fig.3-figure supplement 1) is interesting because it suggests that BF neurons can release ACh alone without GABA as a co-transmitter, unlike what has been reported in deep OB layers (Case, Burton et al. 2017). To confirm this hypothesis, I selectively targeted the expression of ChR2-eYFP in cholinergic neurons using virus injection in the HDB/MCPO of ChAT-cre mice (Fig.4A). Choline acetyltransferase (ChAT) immuno-detection on brain sections from these mice (thereafter called ChAT mice) confirmed the expression of ChR2 in neurons expressing endogenous ChAT (Fig.4B). ChR2 expression, as indicated by eYFP labeling in the HDB/MCPO, was almost exclusively found in cholinergic neurons (89% of the cells positive for eYFP were also ChAT+, 440 double+ cells from a total of 495 eYFP+ cells in coronal slices from 4 mice) with an infection rate of 34% (440 double+ cells from a total of 1305 ChAT+ cells). Consistent with previous studies (Kasa et al. 1995, Rothermel et al. 2014, Smith et al. 2015, Hamamoto et al. 2017), eYFP-labeled cholinergic axons arising from BF cholinergic neurons densely innervated the OB and were abundant in the glomerular layer (Fig.4C).

**Figure 4:**
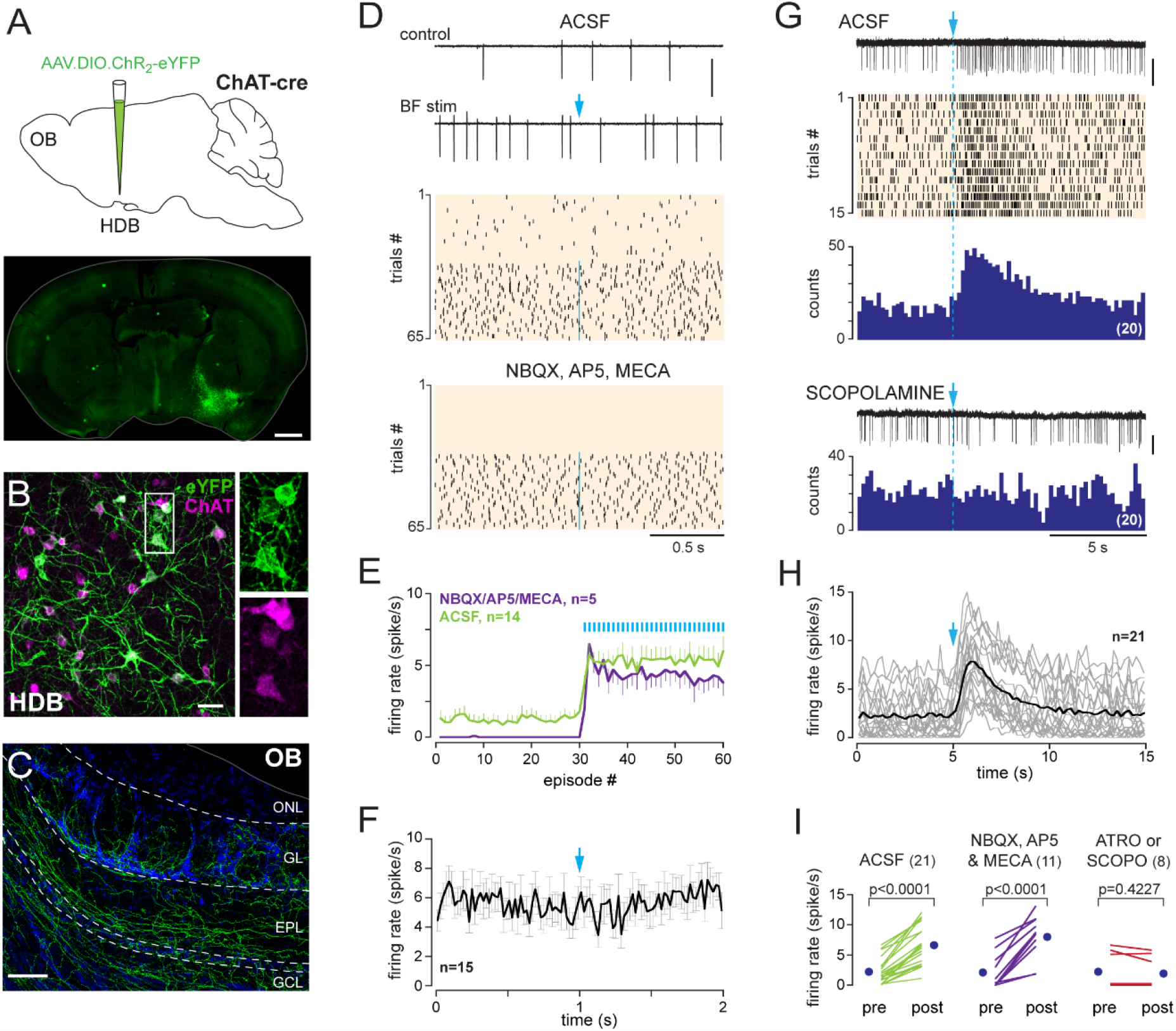
BF cholinergic neurons do not release GABA onto PG cells. **A**, Schematic of the virus injection in ChAT-cre mice and coronal section, at about Bregma -0.1 mm, 20 days after injection. **B**, ChR2-eYFP expression (green) in ChAT-expressing cholinergic neurons (magenta) in the BF. Scale bar 50 µm. Right panels, zoom on the boxed region. **C**, ChR2-eYFP-expressing axons in the OB. Scale bar 100 µm. DAPI staining (blue) delimits layers (ONL: olfactory nerve layer; GL: glomerular layer; EPL: external plexiform layer; GCL: granule cell layer). **D**, Top, two representative traces and raster plot of spiking activity in control condition and when BF axons were photostimulated (blue arrow). Photostimulation started at episode 31 (1 flash/episode, blue line). Bottom, raster plot for the same cell, same experiment in the presence of blockers. **E**, Average firing rate per episode (2 s each) in ACSF (green) or in the presence of blockers (violet). Low frequency photostimulation (0.5 Hz) started at episode 31. **F**, Average firing rate per bin and per episode for 15 cells while BF axons were photostimulated once per episode. BF axons stimulation did not inhibit firing. **G**, representative recording, raster plot and PSTH (20 consecutive trials, bin 200 ms) of spiking in ACSF (top). BF axons photostimulation at blue arrow and dotted line. Scopolamine (10 µM) blocked the evoked excitation (bottom). **H**, Average firing rate per bin and per 15 s-long episode for 21 cells. Each grey line corresponds to a cell, the black line indicates the ensemble average. **I**, Firing rate before (pre) and after (post) photostimulation of BF axons in ACSF (green), in the presence of NBQX, D-AP5 and mecamylamine (violet) or in the presence of atropine (n=4) or scopolamine (n=4)(red). Each line indicates a cell, blue circles are the mean. Paired t-test or Wilcoxon signed-rank sum test (for atro/scopo).

In OB slices from ChAT mice, low frequency photostimulation at 0.5 Hz caused a significant increase in baseline spike frequency in 7% of the PG cells tested (n=23/350). Importantly, this was the only kind of response observed in LCA recordings. As in dlx5/6 mice, responsive cells had a low baseline spiking activity in control condition (range 0-6 Hz, mean 1.2 ± 1.6 Hz, n=14) and switched to a higher frequency firing mode during the 0.5 Hz stimulations in ACSF (mean frequency 5.3 ± 2.7 Hz, n=14) as well as in the presence of NBQX, D-AP5 and mecamylamine (n=5, Fig.4D and 4E). In these cells, a single flash also evoked a robust long-lasting excitation that persisted in the presence of NBQX, D-AP5 and mecamylamine, had a similar time course as in dlx5/6 mice and was blocked by scopolamine or atropine (Fig.4G-I). However, in sharp contrast with the dual inhibition-excitation response in dlx5/6 mice, responses were only excitatory and lacked the transient GABA-mediated post-stimulus spiking inhibition (Fig. 4F and 4H). Hence, these results demonstrate that cholinergic axons mediating a muscarinic excitation at PG cells do not release GABA, implying that separate populations of BF neurons mediate the dual GABAergic-muscarinic responses in PG cells.

### Endogenous ACh release elicits a slow muscarinic EPSC in a specific subtype of PG cell

In our previous report, we did not detect any BF-evoked muscarinic EPSC in whole-cell recordings from various PG cell subtypes (Sanz Diez et al. 2019). To understand the mechanism underlying photo-evoked BF excitation, I re-examined this question using a slightly different K-gluconate-based internal solution supplemented with phosphocreatine to improve ATP supply. PG cells receiving a muscarinic input were first identified using LCA spike recording and then whole-cell (WC) patched with a pipette filled with the new internal solution (n=5 cells in ChAT mice and n=12 in dlx5/6 mice). In dlx5/6 mice, photostimulation of the BF axons evoked a biphasic response initiated by an outward IPSC followed by a slow rising inward current in 7/12 cells (Fig.5A). The slow EPSC was undetectable in 5 other cells. In ChAT mice, BF axons photostimulation induced a slow inward EPSC in 5/5 cells. Consistent with the LCA experiments, the slow inward current was not preceded by an outward IPSC, further confirming that cholinergic axons do not release GABA on these PG cells (Fig.5D). In both transgenic lines, photo-evoked muscarinic EPSCs recorded at negative holding potentials (from -30 to -60 mV) were small and often at the limit of detection (max 10 pA, mean 5 ± 3 pA, data obtained in ChAT and dlx5/6 mice pooled together). In current-clamp, at membrane potential close to the resting potential, photo-evoked EPSPs did not exceed 5 mV (mean 3.1 ± 1.1 mV, 2 cells in ChAT mice and 3 cells in dlx5/6 mice)(Fig.5A and 5D). Unfortunately, muscarinic EPSC/EPSP ran down in few minutes, precluding any further characterization.

**Figure 5:**
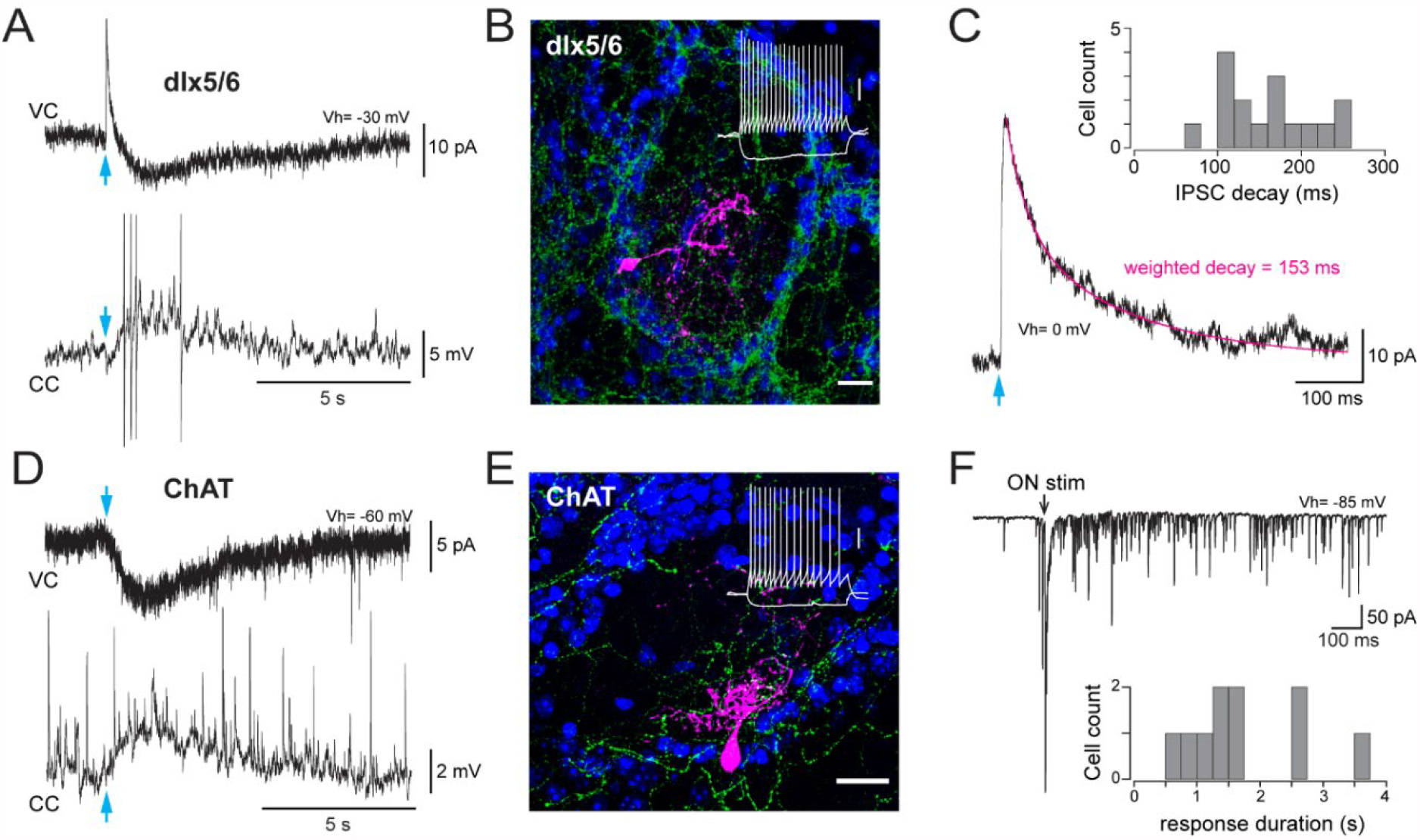
BF cholinergic inputs produce a slow muscarinic EPSC in a specific PG cell subtype. **A**, Biphasic GABAergic-muscarinic voltage-clamp (VC) and current-clamp (CC) whole-cell responses to a single photostimulation (blue arrow) of BF axons in a cell from a dlx5/6 mouse. Responses recorded in the presence of NBQX, D-AP5 and mecamylamine. The current is an average of 8 consecutive sweeps, the voltage response is a single trace eliciting 4 spikes (truncated for display). **B**, Morphology of a biocytin-filled PG cell in which photostimulation of BF axons (eYFP-positive, green) produced a dual GABA-ACh response in a dlx5/6 mouse. DAPI staining (blue) shows the outline of the glomerulus. Scale bar 20 µm. Inset: voltage responses of this cell to depolarizing and hyperpolarising current steps (20 pA, 500 ms). Scale bar 20 mV. **C**, Photo-evoked GABAergic IPSC recorded in the same PG cell as in A at a holding potential of 0 mV. Average of 10 sweeps. The decay was best fitted with two exponentials (magenta) with a weighted decay time constant of 153 ms. Inset: distribution histogram of the decay time constants of photo-evoked GABAergic IPSCs in PG cells with a mixed GABA/ACh response in dlx5/6 mice. **D**, Photo -evoked muscarinic EPSC (VC) and EPSP (CC) recorded in ACSF in a PG cell from a ChAT mouse. Average of 6 consecutive sweeps for the EPSC, single trace for the EPSP. **E**, Morphology of a biocytin-filled cell that responded to the photostimulation of BF cholinergic axons with a muscarinic excitation in a ChAT mouse. Scale bar 20 µm. Blue: DAPI; green: eYFP positive BF axons. Inset: membrane voltage responses of this cell to the injection of current steps (−20/+35 pA, 500 ms). Scale bar 20 mV. **F**, Long-lasting barrage of EPSCs evoked by the electrical stimulation of the olfactory nerves (black arrow, 0.1 ms/100 µA) in the cell shown in E. The distribution histogram shows the duration of the ON-evoked response elicited in 10 cells receiving a muscarinic excitation (7 cells from dlx5/6 mice and 3 cells from ChAT mice).

Intrinsic, synaptic and morphological properties of the responsive PG cells were also examined. In addition to the 17 WC-recorded cells from ChAT and dlx5/6 mice, the dataset included WC recordings from previous experiments in dlx5/6 mice in which a photo-evoked muscarinic excitation was noticed in the cell-attached configuration (n=9). The morphology of each cell was assessed at the end of the recording by visual inspection of the dye present in the internal solution. Moreover, six cells filled with biocytin were successfully recovered for post-hoc morphological reconstruction. All had the typical morphology of PG cells. Their soma was small, ovoid or round, with no apparent axon and projected small thin dendrites within a single glomerulus (Fig.5B and 5E). On average, their electrical membrane resistance was 1061 ± 524 MΩ. They all fired regularly at up to 100 Hz with overshooting action potentials in response to depolarizing current steps (Fig.5B and 5E). Another striking hallmark of this PG cell subtype was their prolonged multi-synaptic response to a single electrical stimulation of the olfactory nerves (ON). This response had no monosynaptic component and consisted of a barrage of fast EPSCs often lasting several hundreds of ms (average 1760 ± 927 ms, n=10)(Fig.5F). Finally, in dlx5/6 mice, photo-evoked BF IPSCs (amplitude range 10-186 pA, mean 62 ± 40 pA) were particularly slow (average decay time constant 162 ± 55 ms, range 69-258 ms, n=16)(Fig.5C). Together, these homogeneous intrinsic and synaptic properties unambiguously identify a specific subtype of type 2 PG cells as the principal and perhaps unique synaptic target of cholinergic BF axons among PG cells. I will refer to this subclass as type 2.3 PG cells to differentiate it from the two previously described classes of type 2 PG cells, i.e. CR-expressing PG cells (here referred as type 2.1)(Fogli-Isepe et al., 2016; Benito et al., 2019) and PG cells labeled in the Kv3.1-eYFP mouse that include calbindin (CB)-positive and CB-negative cells (here referred as type 2.2)(Najac et al., 2015).

To confirm that type 2.3 PG cells were the only PG cell target of BF cholinergic neurons, I examined whole-cell responses evoked by a single flash in other types of PG cells in ChAT mice. Photo-evoked responses were recorded in voltage-clamp at 0 mV and at negative holding potential in order to detect possible monosynaptic GABAergic, nicotinic or muscarinic responses. Membrane properties and ON-evoked responses were examined when possible to identify each cell type. Photostimulation of the BF fibers did not evoke any response in type 1 PG cells (n=4), type 2.1 PG cells (n=21) and type 2.2 PG cells (n=17). 12 additional PG cells that could not be firmly classified did not respond either. Only one PG cell, that could not be unambiguously classified, responded with a fast IPSC, a rare event that possibly reflected unspecific expression of ChR2 in BF GABAergic neurons. Together, these results suggest that BF cholinergic axons innervating the glomerular layer of the bulb do not release GABA and have a unique and selective synaptic target among PG cells.

### M1 mAChR mediates the muscarinic excitation

Next, I determined what muscarinic receptor mediates the cholinergic response and what downstream mechanism produces the excitation. There are five types of mAChRs and three of those (M1, M3 and M5) are coupled to excitatory G_q/11_ proteins that activate PLCβ, causing hydrolysis of PIP2 into DAG and IP3. Activation of these receptors most often depolarizes neurons. M1 receptors are widely expressed in the OB (Le Jeune et al. 1995) and their blockage impairs olfactory-evoked fear learning (Ross et al. 2019) making them a likely candidate. To test this hypothesis, I examined the effects of pirenzepine (1-2 µM), a selective antagonist of M1 receptors, on BF-induced muscarinic excitation in both ChAT (n=3 cells) and dlx5/6 mice (n=7). Experiments were done using LCA recording in the presence of glutamate and nicotinic receptor antagonists. Pirenzepine fully blocked the excitation evoked by light stimulation of the BF fibers in 6 cells and reduced its strength in 4 cells (Fig.6A and 6B). On average, post stimulation spike frequency was reduced 4-fold by pirenzepine (control: 10 ± 3.4 Hz; pirenzepine: 2.6 ± 3.3 Hz, p<0.0001, data from ChAT and dlx5/6 mice pooled together, Fig.6B). These results support the idea that M1 receptors mediate the muscarinic excitation of type 2.3 PG cells.

**Figure 6:**
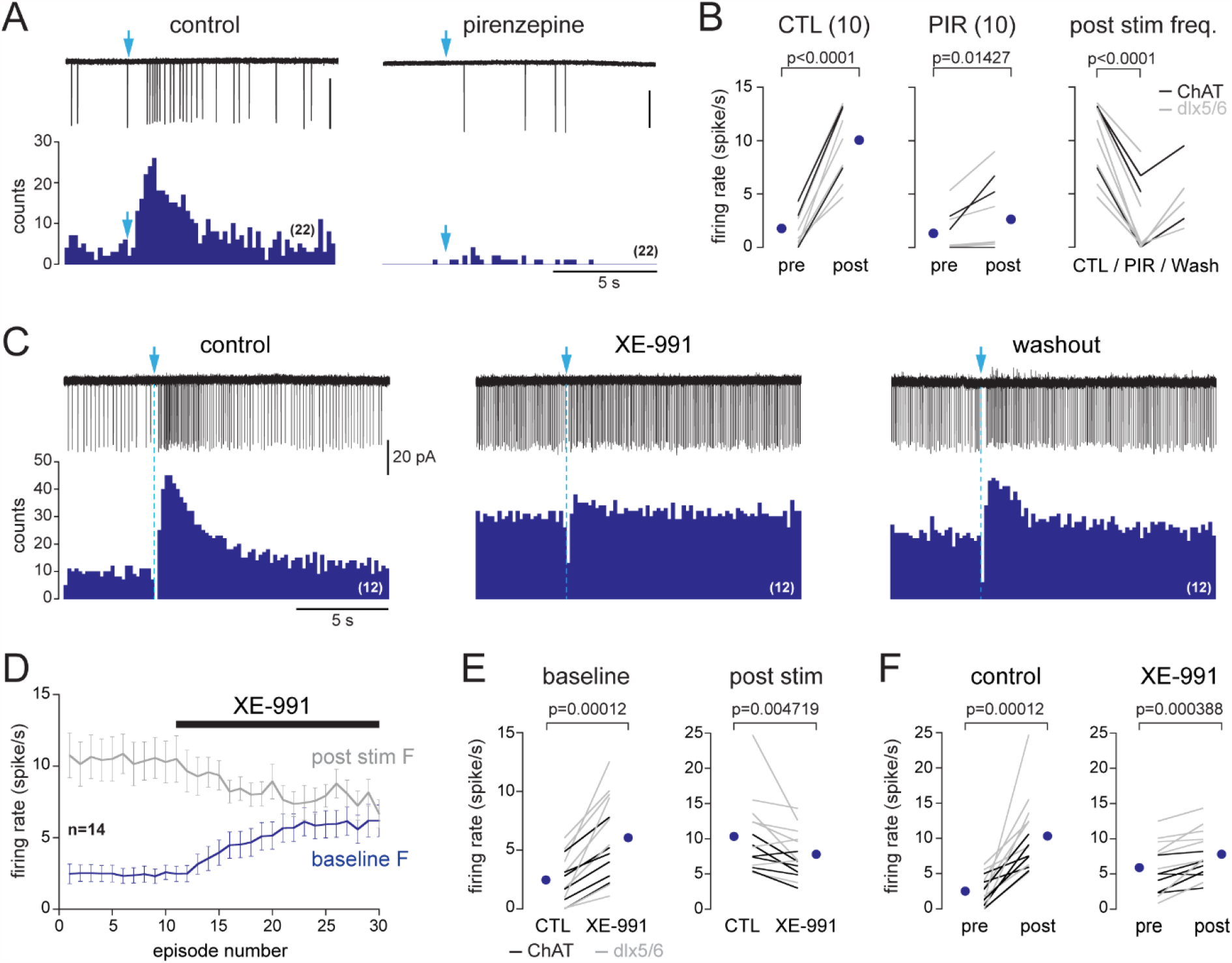
M1 mAChRs mediate BF-evoked excitation by closing M channels. **A**, Photo-evoked LCA responses and cumulative PSTH (over 22 consecutive sweeps, 200 ms/bin) recorded in a PG cell from a dlx5/6 mouse in control condition (left) and in the presence of the M1 mAChR antagonist pirenzepine (2 µM). Scale bar for traces 50 pA. **B**, Summary graphs. Firing rate before (pre) and after (post) photostimulation of BF fibers in control condition (left, paired t-test) and in the presence of pirenzepine (PIR, 1 or 2 µM, paired Wilcoxon signed-rank sum test). Right graph shows that pirenzepine decreased BF-evoked excitation in every cell tested (paired t-test). Partial washout was obtained in 5 cells. Cells were recorded in ChAT mice (n=3, black lines) and in dlx5/6 mice (n=7, grey lines). **C**, Photo-evoked LCA responses and cumulative PSTH recorded in a PG cell from a dlx5/6 mouse showing the effects of the M-channel blocker XE-991 (10 µM) on spiking frequency. BF fibers were photostimulated with a single flash (blue arrow and dotted line). **D**, XE-991 increased baseline spiking rate (blue line, measured during a 15 s time period preceding the flash) and decreased post stimulus spike frequency (grey line). Average from 14 cells (8 in dlx5/6 mice, 6 in ChAT mice). Each episode was 30 s long. **E**, summary graph showing the two effects of XE-991 on each cell. Paired Wilcoxon signed-rank sum tests. **F**, Firing rate before (pre) and after (post) photostimulation of BF fibers in control condition (left, paired Wilcoxon signed-rank sum test) and in the presence of XE-991 (t-test). Experiments were all done in the presence of NBQX (10 µM), D-AP5 (50 µM) and mecamylamine (50 µM). Means are the blue circles.

M1 receptors classically suppress the M current, a slow voltage-activated potassium current mediated by M-channels and active at resting membrane potential (Suh and Hille 2008, Brown and Passmore 2009). The fast run-down of the BF-evoked muscarinic EPSC in whole-cell recordings as well as its strong dependence on intracellular ATP are consistent with this downstream mechanism. Cell dialysis is indeed often deleterious for M-currents because M-channels opening depends on PIP2 binding, a process that is itself highly dependent on intracellular ATP supply for PIP2 phosphorylation (Zhang et al. 2003, Suh and Hille 2008). To investigate whether closure of the M current causes the muscarinic depolarization of type 2.3 PG cells, I applied the selective M-channel antagonist XE-991 (10 µM) in the presence of NBQX, D-AP5 and mecamylamine in LCA. XE-991 had two noticeable effects in type 2.3 PG cells. First, it increased baseline spike frequency in all the cells tested (n=14, 8 in dlx5/6 mice and 6 in ChAT mice, Fig.6C-E) suggesting that M-channels are indeed open at rest in type 2.3 PG cells and hyperpolarize their membrane potential. Second, although XE-991 did not fully block photo-evoked muscarinic excitation in most cells (Fig.6C and in Fig.6F), it reduced the strength of BF-induced excitation (Fig. 6D and 6E). A persistent photo-evoked excitation is expected if the muscarinic EPSP is only partially blocked while the membrane potential is more depolarized compared to control condition. Overall, these data support the hypothesis that M1 mAChRs activation depolarizes type 2.3 PG cells by blocking a potassium M current.

### Muscarinic excitation of type 2.3 PG cells leads to an increase of inhibition in principal neurons

PG cells are classically viewed as a source of intraglomerular inhibition of mitral and tufted cells, the two OB output channels that project to distinct cortical areas. However, the targets and output properties of type 2.3 PG cells have never been specifically investigated. Thus, to evaluate the impact of the muscarinic excitation of type 2.3 PG cells, I recorded IPSCs in mitral cells and in superficial or middle tufted cells (s/mTC) in the presence of NBQX, D-AP5 and mecamylamine (Fig.7A). I also examined inhibition in external tufted cells (eTCs)(Fig. 7A). Although it is still unclear whether eTCs have axonal projections outside the OB like other tufted cells, they play a major role in processing incoming OSN information by coordinating rhythmic activity within each glomerulus and by providing feedforward excitation to various types of neurons including mitral and tufted cells (Hayar et al. 2004, De Saint Jan et al. 2009, Najac et al. 2011, 2015).

**Figure 7:**
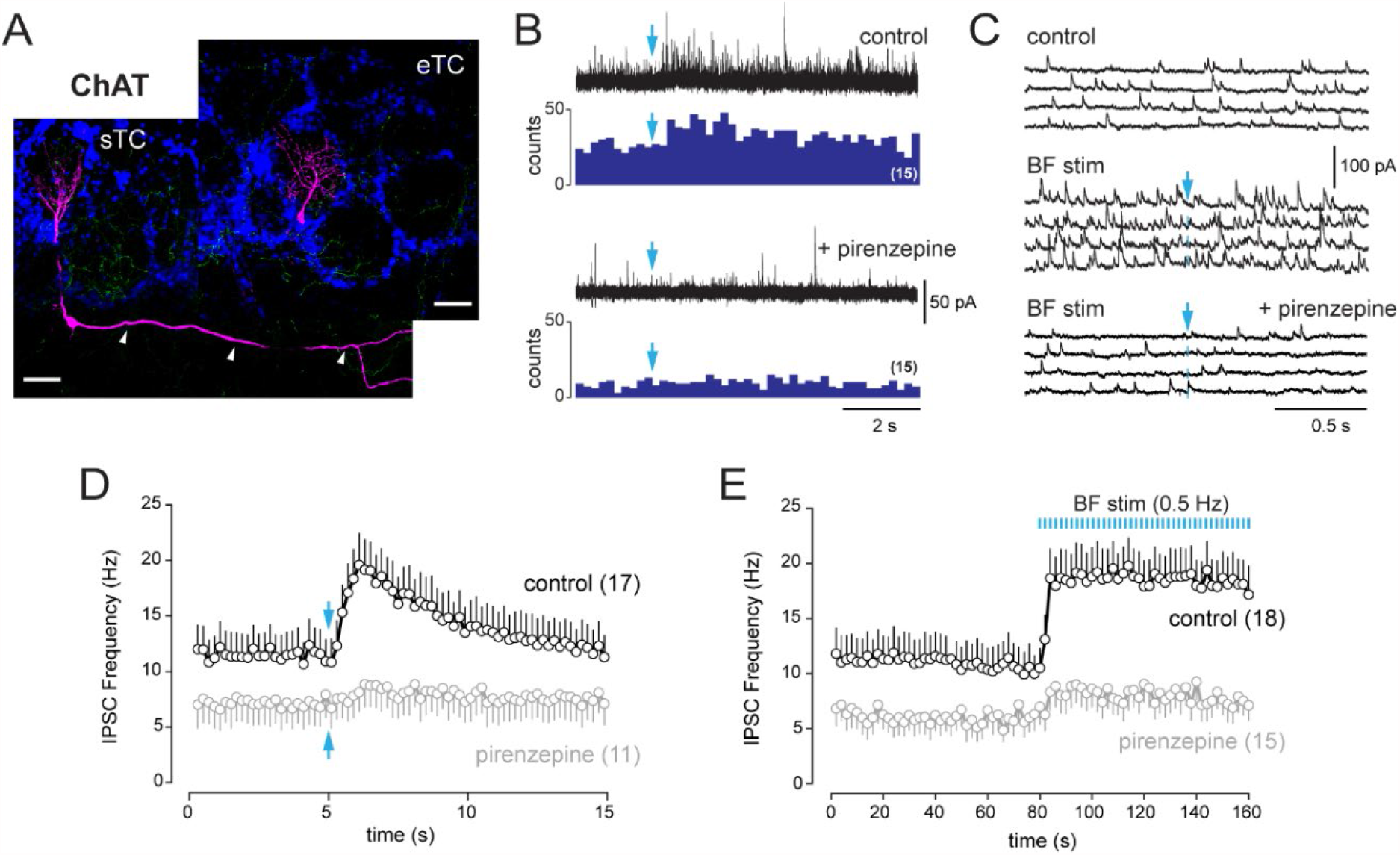
BF Muscarinic inputs lead to an increase of inhibition in principal neurons. **A**, Morphologies of two biocytin-filled tufted cells recorded in different slices from ChAT mice. The localization of the soma and the presence or not of lateral dendrites in the external plexiform layer (arrowheads) distinguish a sTC (left) and an eTC (right). DAPI staining (blue) shows the outline of glomeruli. ChR2-eYFP-expressing cholinergic fibers are visible in green. Scale bars 50 µm. **B**, Increase of inhibition evoked by a single photostimulation (blue arrow) of the cholinergic axons in a sTC (top). Pirenzepine (2 µM) blocked this response (bottom). 5 traces are superimposed in each conditions. PSTH show the cumulative number of IPSCs/bin (200 ms) across 15 consecutive trials. **C**, Four consecutive traces of spontaneous IPSCs recorded in an eTC in control conditions (top) and during low frequency photostimulation of the cholinergic BF fibers (1 flash every 2 s at blue arrow, middle). Pirenzepine (2 µM) blocked the increase of IPSC frequency evoked by the photostimulations (bottom). **D**, Average IPSC frequency per 200 ms bin and per episode for 17 tufted cells (8 s/mTC and 9 eTC). BF axons were photostimulated once (blue arrow). Pirenzepine was tested on 11/17 cells and reduced photo-evoked increase of IPSCs. **E**, average IPSC frequency per episode (2 s) for 18 cells (10 s/mTC and 8 eTC). Photostimulation of the cholinergic fibers at 0.5 Hz rapidly and persistently increased IPSC frequency. Pirenzepine was tested in 15/18 cells. Experiments were all done in ChAT mice in the presence of NBQX, D-AP5 and mecamylamine. Individual data points are shown in Figure 7-Figure supplement 1.

Photostimulation of the cholinergic axons significantly increased IPSC frequency in eTCs (n=9) and in s/mTCs (n=10) (Fig.7) whereas no response was found in mitral cells (n=17). However, three of the recorded mitral cells had severed apical dendrites and most of the intact cells (10/14) projected in glomeruli located deep within the slice, where light stimulation may be less efficient. Baseline IPSC frequency greatly varied across cells and was on average smaller in s/mTC (7.0 ± 5.1 Hz) compared to eTC (15.7 ± 10 Hz, p=0.0285, t-test). Yet, IPSCs increased in similar proportion and with comparable time course in both cell types after photostimulation (Figure 7-figure supplement 1). Data were thus pooled in Fig.7D and 7E. Thus, low frequency photostimulation (1 flash every 2 s) led to a rapid and persistent increase in IPSC frequency in 8/10 eTC and in 10/23 s/mTC. This response was blocked (n=8) or reduced (n=3) when the experiment was repeated in the presence of 2 µM pirenzepine (Fig.7C and 7E). Pirenzepine had little effect in 4/15 cells. A single photostimulation also transiently increased IPSC frequency in eTC (n=9) and s/mTC (n=8)(Fig.7B and 7D). Addition of pirenzepine attenuated this response in 10/11 cells. Together, these results are consistent with an increased inhibition of tufted cells by type 2.3 PG cells following their excitation by muscarinic inputs. This increased inhibition could lead to profound changes on the activity and output of the OB network.

### BF GABAergic inputs inhibit type 2.2 PG cells

Beside type 2.3 PG cells described above, a second group of cells in dlx5/6 mice responded to the optogenetic stimulation of the BF axons with a transient block of spiking immediately after the photostimulation (Fig.8). However, unlike type 2.3 PG cells, they did not show any evidence of parallel cholinergic excitation. Baseline firing activity did not rapidly increase when BF fibers were photostimulated at 0.5 Hz and a single flash did not elicit a long-lasting mAChR-evoked increase in firing (Fig.8-figure supplement 1). In these cells, basal firing frequency was, on average, higher than in cells responding with a dual inhibition-excitation (mean 7.8 ± 5.6 Hz, n=21, p=0.00058, Mann-Whitney Rank Sum Test). However, the nature of this activity (single spike vs. burst of spikes) and its frequency (range 2-21 Hz) varied greatly across cells (Fig.8-figure supplement 2). Spontaneous spiking was also diversely affected by NBQX, D-AP5 and mecamylamine, being totally blocked in some cells (n=3) and not affected in others (n=9). In the latter, BF-induced inhibition persisted in the presence of the blockers (n=9) and was blocked by gabazine (Fig.8B, n=7) indicating that it was caused by a direct GABAergic BF input. The duration of the BF-induced inhibition also varied across cells. In about half of the cells, the BF input induced a shorter (<200 ms) inhibition than in type 2.3 PG cells whereas inhibition was longer and similar as in type 2.3 PG cells in others (Fig.8C). However, it is noteworthy that cells with a prolonged inhibition often fired at low rate or fired irregularly with bursts of spikes (Fig.8-figure supplement 2). In these cells, the duration of BF-induced inhibition, as measured as the mean delay between the flash and the first spike after the flash, varied across sweeps making this estimate imprecise (Fig.8-figure supplement 2A).

**Figure 8:**
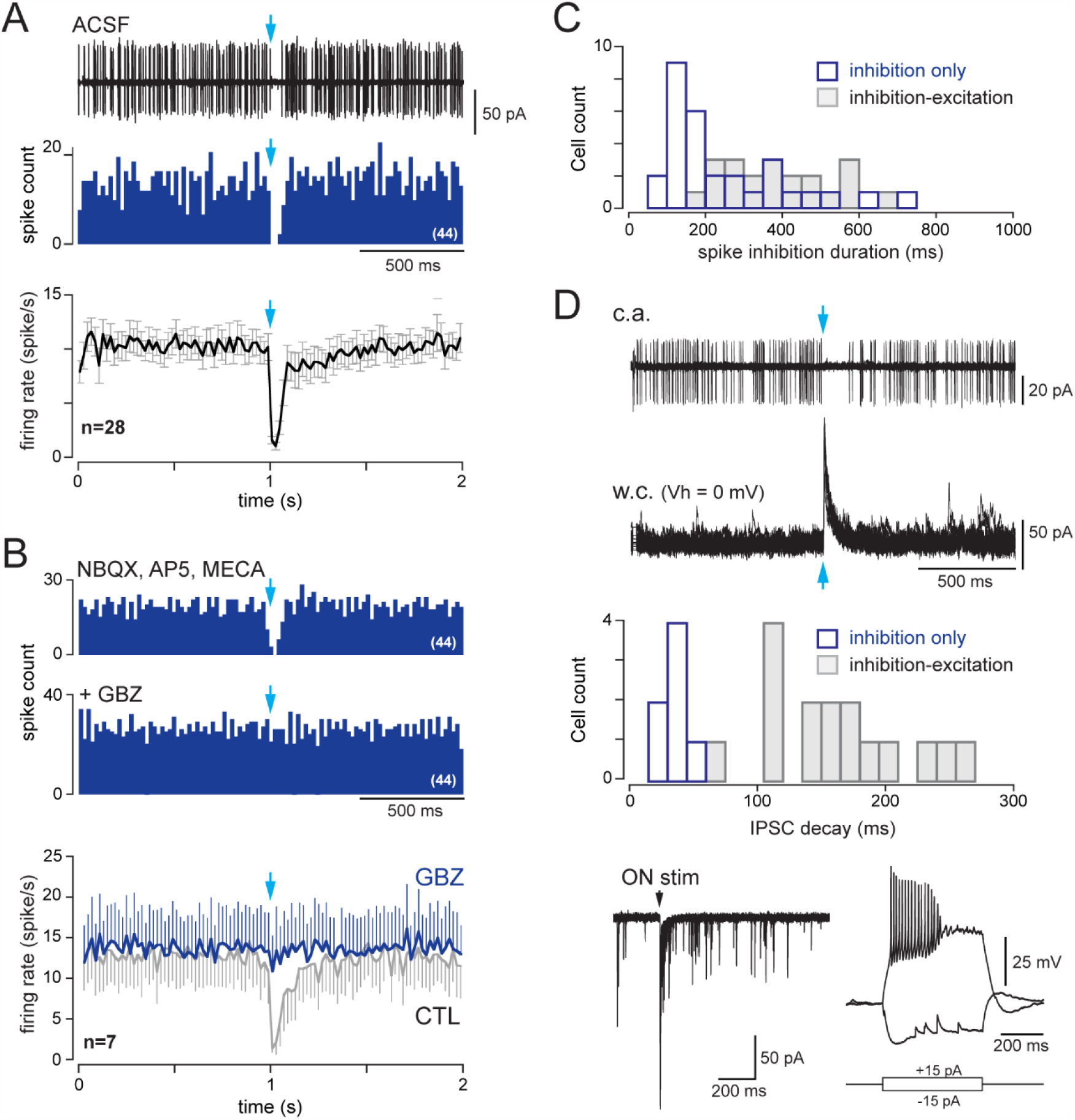
type 2.2 PG cells are inhibited by the BF GABAergic input. **A**, 10 superimposed LCA traces and the corresponding PSTH for 44 consecutive trials (bin 20 ms) in a cell from a dlx5/6 mouse. Photostimulation of the BF fibers (blue arrow) transiently blocked spiking. Bottom, average firing frequency per bin (20 ms) and per episode for 28 cells that were inhibited by the BF input, without evidence of parallel cholinergic excitation (Figure 8-figure supplement 1). **B**, BF-induced spiking inhibition persisted in the presence of NBQX, AP5 and mecamylamine but was blocked by gabazine. The two PSTH are from the same cell as in A (bin 20 ms). Bottom, average firing rate per bin (20 ms) and per episode for 7 cells in control conditions (grey line, 6 cells in the presence of blockers, 1 cell in ACSF) and when GBZ (5 µM) was added (blue). **C**, Duration of the post-stimulus spiking inhibition in cells with a BF-induced inhibitory response (white) vs. in cells with a dual inhibition-excitation response (grey). See also Figure 8-figure supplement 2 for caveats in these measurements. **D**, Whole-cell characterization of a cell with an inhibitory response. Top, BF impact on firing (cell-attached recording, 38 episodes are superimposed) and BF-evoked IPSC (whole-cell recording, 15 superimposed episodes). The histogram compares the decay time constants of photo-evoked GABAergic IPSCs in 7 PG cells that were only inhibited (white bars) vs. in cells with a mixed GABA/ACh response (grey bars). Bottom, ON-evoked EPSC (left, 4 superimposed traces) and current-clamp voltage responses to current steps.

Cells with BF-induced spiking inhibition and no evidence for cholinergic excitation in the cell-attached configuration were subsequently characterized in the whole-cell mode (n=7, Fig.8D). Their electrical membrane resistance was 1119±789 mOhm. BF axons stimulation evoked large IPSCs (amplitude range 69-464 pA, mean 170±144 pA) with an average decay time constant of 37±9 ms, that was significantly shorter than in type 2.3 PG cells (p<0.0001, Mann-Whitney Rank Sum Test). Three cells in which ON-evoked responses were recorded responded with a short pluri-synaptic burst of EPSCs (duration <150 ms). Firing properties were more heterogeneous. Injection of a depolarizing current step evoked a burst of action potentials followed by a plateau (n=3 cells, as in the example shown in Fig.8E) or sustained firing of action potentials (n=3, not shown). In 5/6 cells, injection of an hyperpolarizing step caused a voltage sag suggesting the activation of an Ih current. All together, these properties match well with those of type 2.2 PG cells, a mixed population of CB-positive and CB-negative PG cells labeled in the Kv3.1-eYFP mouse (Najac et al. 2015). This suggests that type 2.2 PG cells only receive an inhibitory GABAergic input from the BF.

### GABA is excitatory in a fraction of CR-expressing type 2.1 PG cells

Finally, photo-evoked BF inputs elicited a single spike (Fig.9A) or, more rarely, a doublet in a small number of cells in dlx5/6 mice (Fig.9C). These excitatory responses were not seen in ChAT mice, persisted in the presence of NBQX, D-AP5 and mecamylamine (n=6) and were blocked by gabazine (n=4, Fig.9B and 9D) suggesting that they were caused by a direct excitatory BF GABAergic input. It would not be surprising if GABA was depolarizing in CR-expressing PG cells because they retain many properties of newborn immature neurons (Benito et al. 2018). Consistent with this hypothesis, most of the cells with an excitatory response (26/29) had no or little (<1 Hz) spontaneous firing activity, similar as CR+ PG cells (Fogli Iseppe et al. 2016, Benito et al. 2018). However, photo-evoked action potentials occurred within a short delay after the flash, as expected for synaptically-driven spikes, in only 15 cells (average spike timing 13.4 ± 9.9 ms, Fig.9A and 9E). In the other cells, the average delay was longer (329 ± 191 ms) and more variable (n=14, Fig.9C and 9E) suggesting that distinct mechanisms drive the two types of response. Strikingly, BF synaptic inputs also evoked a small but detectable gabazine-sensitive capacitive current that was inward in 14/15 cells responding with early spikes (Fig.9A-B, 9E) whereas it was outward in all the cells responding with delayed spikes (Fig.9C-D, 9E). Thus, a likely explanation of the data is that a depolarizing GABAergic input directly triggered early spikes whereas delayed spikes could be induced by rebound depolarization following an hyperpolarizing GABAergic IPSP (as often seen in CR+ PG cells)(Benito et al. 2018). Eight cells responding with early action potentials in the cell-attached mode (average timing 20 ± 19 ms) were subsequently characterized in the whole-cell configuration. All of them had properties consistent with those of CR+ PG cells, i.e. a large input resistance (6 ± 3.1 GΩ), a characteristic voltage response to depolarizing current steps and a fast and large photo-evoked BF IPSC (amplitude range 133-1300 pA, mean 370 ± 139 pA; weighted decay time constant 7.7 ± 3.6 ms)(Fig.9G).

**Figure 9 :**
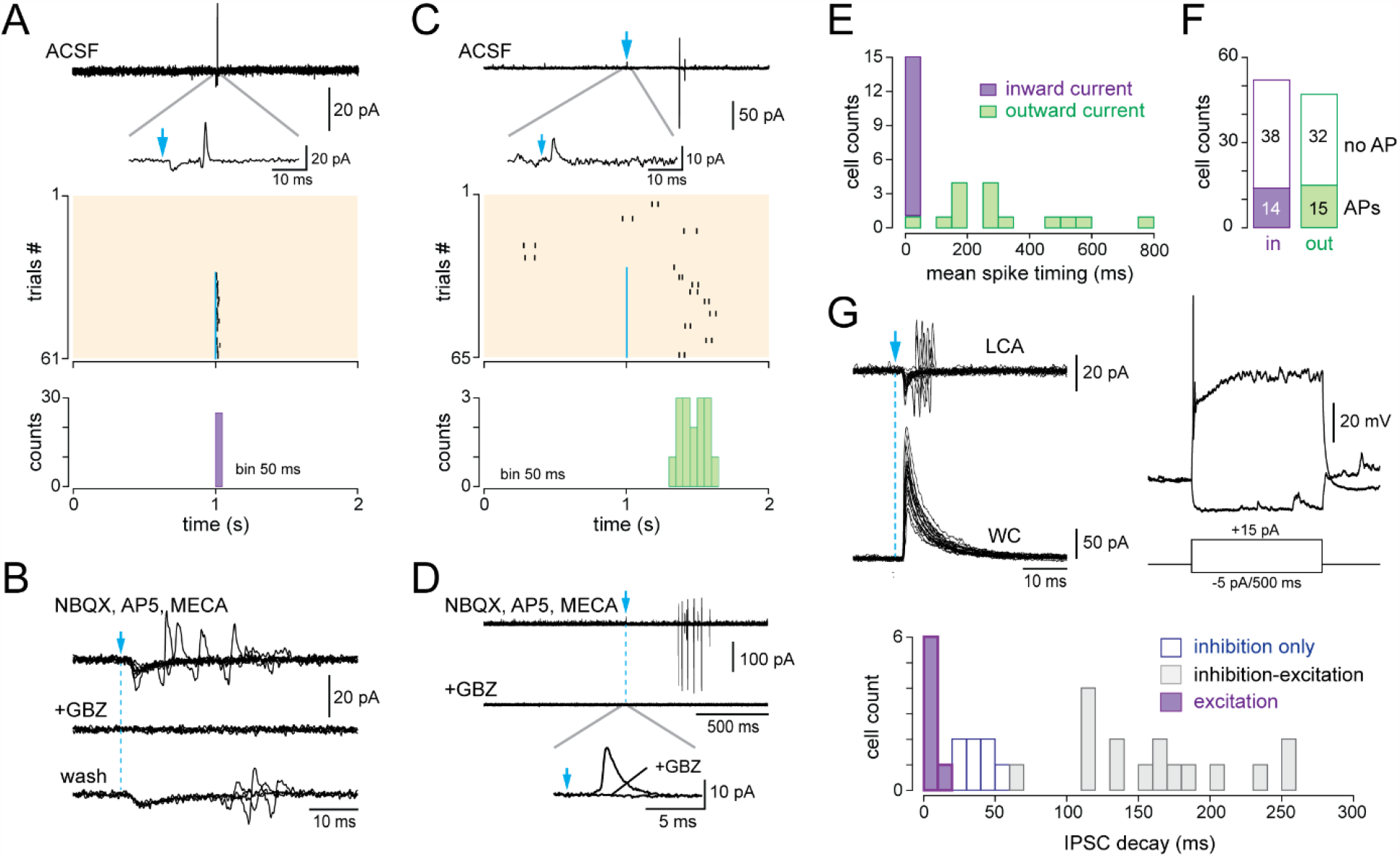
BF-evoked GABAergic excitation in a fraction of type 2.1 PG cells. **A**, example of a BF-evoked excitatory response in a PG cell. One representative LCA recording episode (duration 2 s) is shown on top. Photostimulation of the BF fibers (blue arrow) induced an inward current followed by a single spike (inset). Each tick is a spike in the raster plot of these cell responses (stimulation: episodes 31-61). Bottom, corresponding PSTH. Note the temporal precision of the evoked spikes. Bin size 50 ms. **B**, Same cell as in A. The evoked inward current and the evoked spike both persisted in NBQX, D-AP5 and mecamylamine but were blocked by gabazine. At least four traces are superimposed in each condition. **C**, Another example of PG cell responding to the photostimulation with action potentials. Photostimulation (blue arrow, episodes 31-65 in the raster plot) induced an outward current (inset) and, in some trials, a delayed doublet of spikes. Bottom, corresponding cumulative PSTH (bin size 50 ms). **D**, Same cell as in C. Gabazine blocked both the evoked outward current and the evoked spikes. **E**, Distribution histogram of the average spike timing in each cell responding with spikes. Cells in which the stimulation evoked an inward capacitive current are in violet, cells in which the stimulation evoked an outward current are in green. **F**, Total number of cells responding with an inward (violet) or an outward (green) capacitive current followed or not with evoked-spikes. **G**, Whole-cell caracterization of a cell excited by the BF input. Left, BF-evoked spike response (LCA recording, top, 20 consecutive episodes are superimposed) and BF-evoked IPSC (whole-cell recording, 20 consecutive responses). Right: current-clamp voltage responses to current steps in the same cell. The distribution histogram compares the decay time constants of photo-evoked GABAergic IPSCs in the 7 PG cells that were excited (violet bars) vs. in type 2.2 cells with an inhibitory response (white) and in type 2.3 cells with a mixed GABA/ACh response (grey bars).

The BF input induced a detectable capacitive current, but no spike, in 70 additional cells, all with no or little spontaneous activity (these cells were classified as non-responsive in Fig.1C). BF-evoked capacitive current were inward in 54% of these cells (n=38/70) and outward in the other cells (n=32, Fig.9F). Eight of these cells were characterized with whole-cell recording and all displayed the typical intrinsic and synaptic properties of CR+ PG cells including a high membrane resistance (3±1.2 GΩ) and large and fast BF IPSCs (amplitude: 284±107 pA; decay 12.2±2.9 ms). All together, this analysis suggests that the BF GABAergic input is depolarizing in as much as half of the CR-expressing type 2.1 PG cells and drives spiking in a minority of them.

## Discussion

This studies shows that OB-projecting BF neurons have diverse impacts on PG cells. GABAergic inputs potently block the discharge of type 2.2 and type 2.3 PG cells with a target-specific time course. In contrast, GABA release is excitatory and eventually triggers action potentials in a fraction of type 2.1 PG cells. Data also reveal that BF cholinergic fibers strongly and exclusively excite type 2.3 PG cells. Thus, intraglomerular inhibition of principal neurons mediated by PG cells can be modulated in various ways by multiple BF pathways that potentially regulate olfactory processing in a context and behavior-specific manner.

### A previously ignored cholinergic pathway

The main finding of this study is that endogenous phasic ACh release from BF cholinergic neurons selectively evokes a remarkably strong and reliable muscarinic excitation in a previously overlooked PG cell subtype, which are referred to as type 2.3 PG cells. This novel cholinergic pathway concerns a small population of neurons lacking a molecular marker and evokes small muscarinic EPSP/EPSCs that rapidly runs down in whole-cell recording, explaining why it has been missed until now. Here, I provide pharmacological evidence suggesting that the slow cholinergic response is mediated by M1 mAChRs that suppress a M current. This downstream mechanism classically washes out quickly and future experiments using perforated patch-clamp experiments will be necessary to confirm this hypothesis.

This pathway adds to other mAChR-dependent mechanisms capable of increasing tonic inhibition in mitral and tufted cells. Activation of M1 receptors increases the excitability of granule cells, the most abundant interneurons in the OB, by potentiating current-evoked afterdepolarization (Pressler et al. 2007). mAChR activation also directly enhances transmitter release at reciprocal dendrodendritic synapses between mitral and granule cells (Castillo et al. 1999, Ghatpande and Gelperin 2009) or between juxtaglomerular interneurons and mitral/tufted cells (Liu et al. 2015). However, there is yet no evidence that endogenous ACh can recruit these previously described pathways. In the previous studies, ACh or cholinergic agonists were exogenously applied on slices. This results in prolonged and uniform activation of synaptic and extrasynaptic ACh receptors and induces multiple concomitant effects that may not necessarily be evoked by physiological release of ACh, even in case of strong afferent activity in cholinergic neurons leading to diffuse volume transmission (Unal et al. 2015).

### Physiological implications

Like other sensory systems, olfactory perception is context-dependent. This modulation already takes place in OB circuits, where odor-evoked neural responses depend on reward (Doucette and Restrepo 2008) or on the difficulty of the task (Koldaeva et al. 2019) and are shaped by learning and experience (Martin et al. 2004, Chu et al. 2017, Ross and Fletcher 2018). Whether BF cholinergic innervation of the OB plays a role in context-dependent neuromodulation is suspected, but has never been proven.

In the cortex, transient increased cholinergic signaling signals a transition to a behaviorally important context and adjust neural output to improve task performances (Pinto et al. 2013, Gritton et al. 2016, Kuchibhotla et al. 2017). In pavlovian learning paradigms, BF cholinergic neurons respond with brief and temporally precise burst of activity to reward or aversive stimuli and to conditioning stimuli, including olfactory cues (Hangya et al. 2015, Guo et al. 2019, Crouse et al. 2020, Hanson et al. 2021). This results in fast and precise transient of ACh in target sensory cortical areas (Guo et al. 2019) or in the basolateral amygdala (Crouse et al. 2020). The physiological dynamics of ACh within the OB are unknown but it is tempting to speculate that similar ACh transients are evoked in the OB during olfactory guided aversive or appetitive learning, a behavior that critically depends on M1 mAChRs in the OB (Ross et al. 2019).

I showed that a single stimulation of the cholinergic BF axons, which evokes transient, temporally and spatially precise release of ACh, triggers a long-lasting discharge in type 2.3 PG cells. This target-specific muscarinic response likely involves synaptic or perisynaptic mAChRs and provides support for phasic, spatially restricted cholinergic transmission as opposed to spatially diffuse volume transmission (Sarter and Lustig, 2020). The same stimulus repeated every two seconds, a low frequency stimulation that is insufficiently strong to induce massive diffusion of ACh in the extracellular space, rapidly transforms type 2.3 PG cells that usually fire at low rate into tonically active neurons that fire at high frequency. This, in turn, leads to a rapid and persistent increase of synaptic inhibition in principal neurons, thus potentially driving OB circuits in a different state of activity. This target-specific muscarinic transmission may have widespread circuit implications because single cholinergic axons frequently ramify and innervate multiple glomeruli in different OB areas (Hamamoto et al. 2017). This could ultimately reduce the firing rate in output neurons but could also shape the temporal structure of mitral and tufted cells output at diverse time scales. Inhibitory inputs regulate spike timing and synchrony in mitral and tufted cells (Schoppa 2006, Shao et al. 2012, Najac et al. 2015) and inhibition of eTCs might modulate slow glomerulus-specific coordinated activity (Hayar et al. 2004, De Saint Jan et al. 2009, Najac et al. 2011). Based on previous studies on ACh functions, increased inhibition driven by the muscarinic excitation of type 2.3 PG cells may improve olfactory perception of behaviorally important odorants. Future *in vivo* experiments will be necessary to explore these possibilities. Interestingly, BF GABAergic fibers innervating type 2.3 PG cells potently block their activity and could act as a powerful brake to reverse the cholinergic effects.

I*n vivo*, optogenetic stimulation of the cholinergic axons in the OB of ChAT-cre mice increases mitral and tufted cells spontaneous and odor-evoked firing (Rothermel et al. 2014, Bohm et al. 2020). This result seems at odds with the expected implications of the new muscarinic pathway described in this study. However, photostimulation was strong and sustained in these *in vivo* studies (light continuously on for 10 s). Although it is difficult to compare stimulations *in vivo* and in slices, a prolonged photostimulus could recruit additional cholinergic pathways that need volume transmission to be activated and that have opposite impacts compared with those of type 2.3 PG cells. For instance, tyrosine hydrowylase (TH)-expressing dopaminergic/GABAergic juxtaglomerular neurons express the M2 mAChR (Crespo et al. 2000, Hamamoto et al. 2017). Activation of these receptors inhibits tonically active TH-expressing cells (Pignatelli and Belluzzi 2008) that provide an inhibitory drive to mitral and tufted cells (Whitesell et al. 2013, Liu et al. 2016, Zhou et al. 2020). Thus, understanding the physiological impact on the OB output and network activity of the muscarinic excitation of type 2.3 PG cells will require targeted stimulation that selectively engages this pathway *in vivo*.

### Multiple cell-type specific pathways for BF control of glomerular inhibition

Results of this study also provide new insights into PG cell diversity. Immunohistochemical studies have already demonstrated that the few classical markers commonly used to label PG cells do not label all of them (Panzanelli et al. 2007, Parrish-Aungst et al. 2007, Whitman and Greer 2007). Yet, most functional studies only distinguish type 1 and type 2 PG cells and ignore type 2’s diversity. CR-expressing type 2.1 PG cells are by far the most abundant, representing 40 to 50% of the entire PG cell population. They are predominately generated postnatally and persist in an immature stage in terms of connectivity and membrane properties (Benito et al. 2018). They receive a fast GABAergic input from BF neurons (Sanz Diez et al. 2019). The present data suggest that this input is excitatory in a fraction of them, most likely the most immature, whereas GABA is inhibitory in the other PG cells. The functional impact of this excitation is unclear because there is no evidence that immature CR+ PG cells form output synapses. However, this result explains why GABA appears predominantly excitatory in calcium imaging of unidentified PG cells (Parsa et al. 2015). Type 2.3 PG cells constitute about 20% of the whole PG cell population and are approximately as numerous as type 2.2 PG cells (Sanz Diez et al. 2019). These regular spiking interneurons have pluri-synaptic long-lasting ON-evoked responses and receive remarkably slow BF IPSCs that readily distinguish them from type 2.1 and type 2.2 PG cells. As shown here, their muscarinic input is another selective feature. Although their input and output connections are not firmly established, type 2.3 PG cells are presumably activated by the glutamate released from mitral and tufted cell dendrites and most likely release GABA unselectively onto mitral and tufted cells. Consistent with this idea, a previous study showed that mAChR activation within glomeruli increases IPSCs equally well in mitral and tufted cells (Liu et al. 2015). The present data confirm that type 2.3 PG cells inhibit various classes of tufted cells but I found no evidence that they also inhibit mitral cells. However, this negative result has to be interpreted with caution as it is challenging in slices from adult mice to find mitral cells projecting in surface glomeruli, a technical requirement for optimal LED stimulation of the cholinergic afferents.

The functional implications of PG cells diversity are not known. Cholinergic and GABAergic inputs from the BF may provide physiological tools to manipulate each PG cell subtype selectively in future studies exploring this question. BF GABAergic axons innervating PG cells indeed have target-specific release properties suggesting that they arise from distinct populations (Sanz Diez et al. 2019). Like elsewhere in the brain, BF GABAergic neurons are highly diverse and each cell population makes cell type-specific long-range connections (Do et al. 2016) and plays specific functions. For instance, somatostatin (SOM)- and parvalbumin (PV)-expressing sub-populations have distinct impacts on arousal control (Xu et al. 2015, Anaclet et al. 2018) or on food intake (Zhu et al. 2017). Similarly, muscarinic excitation of type 2.3 PG cells may involve a specific population of BF cholinergic neurons. There are at least two distinct types of BF cholinergic neurons that differ in their firing modes and synchronization properties and that are differently engaged during behaviors (Laszlovszky et al. 2020). This specificity could also rely on connectivity. For instance, BF cholinergic neurons modulating distinct areas are driven by specific combinations of synaptic inputs (Zaborszky et al. 2015, Do et al. 2016, Gielow and Zaborszky 2017, Zheng et al. 2018). The recent discovery of a genetically-defined sub-population of cholinergic neurons that selectively innervates a specific subgroup of deep short axon cells in the OB (Case et al. 2017) also supports the hypothesis of cell-specific innervation of OB interneurons by specific subsets of cholinergic neurons. Hence, each of the BF neuromodulatory pathways innervating PG cells might be independently recruited during specific tasks or internal states.

## Materials and Methods

### Animals and ethical approval

All experiments procedures were approved by the French Ministry and by the local ethic committee for animal experimentation (CREMEAS). Mice were housed in the animal facility with ad libidum access to food and water. Adult heterozygous dlx5/6-Cre mice (n=36, 34 females - 2 males, C57BL6/J background) and ChAT-Cre mice (n=44 of either sex, CD1 background) were used in this study.

### Stereotaxic viral injection

3-8 weeks old mice were anesthetized with intraperitoneal injection of Zoletil 50 (TILETAMINE/ZOLAZEPAM, 60-70 mg/kg) and Rompun 2% (xylasine, 18-20 mg/Kg) and placed in a stereotaxic apparatus. Metacam (meloxicam, 2 mg/Kg, SC injection) and Lurocaine + Bupivacaine (2 mg/Kg both, SC, local) were administered prior incision. Mice were craneotomized and a volume of 300-400 nl of AAV9.EF1a.DIO.hChR2(H134R).eYFP.WPRE.hGH was stereotaxically injected in the left hemisphere at 0/+0.2 mm AP, 1.4/1.6 mm ML and 5.4 /5.6 mm DV from bregma. Viruses were purchased from the University of Pennsylvania Viral Vector Core (Addgene # 20298) or from the molecular tools platform at the Centre de recherche CERVO (Québec, Canada). After surgery, Antisedan (atipamezol)(2.5%) was injected IP and mice were rehydrated with 0.5 ml of NaCl 0.9% and placed under a heating lamp. Mice recovered during 2-4 weeks after injection before anatomical or physiological experiments.

### Slice preparation

Mice were killed by cervical dislocation and the OB rapidly removed in ice-cold oxygenated (95% O_2_-5% CO_2_) cutting solution containing (in mM): 83 NaCl, 26.2 NaHCO3, 1 NaH2PO4, 2.5 KCl, 3.3 MgSO4, 0.5 CaCl_2_, 70 sucrose and 22 D-glucose (pH 7.3, osmolarity 300 mOsm/l). Horizontal olfactory bulb slices (300 µm-thick) were cut using a Microm HM 650V vibratome (Microm, Germany) in the same solution; incubated for 30-40 minutes at 34°C; stored at room temperature in a regular artificial cerebrospinal fluid (ACSF) until use. ACSF contained (in mM): 125 NaCl, 25 NaHCO3, 2.5 KCl, 1.25 NaH_2_PO4, 1 MgCl_2_, 2 CaCl_2_ and 25 D-glucose and was continuously bubbled with 95% O_2_-5% CO_2_.

### Electrophysiological recordings

Slices were transferred to a recording chamber and perfused with ACSF at 32-34°C under an upright microscope (SliceScope, Scientifica, Uckfield, UK) with differential interference contrast (DIC) and fluorescence optics. Spontaneous action potential firing activity was monitored using loose cell-attached (LCA) recording (15-100 MΩ seal resistance). LCA recordings were made in the voltage-clamp mode of the amplifier (multiclamp 700B, Molecular Devices, Sunnyvale, CA) with no current injected through the pipette. In these conditions, large and fast membrane potential changes such as action potentials are detected as capacitive currents flowing across the patch capacitance (Barbour and Isope 2000). A regular patch pipette filled with ACSF was used on several successively recorded cells. Whole-cell PG cell recordings were made with glass pipettes (4-7 MΩ) filled with a regular K-gluconate-based internal solution containing (in mM): 135 K-gluconate, 2 MgCl_2_, 0.025 CaCl_2_, 1 EGTA, 4 Na-ATP, 0.5 Na-GTP, 10 Hepes (pH 7.3, 280 mOsm, 15 mV junction potential). The intracellular solution used to record the muscarinic EPSC was adapted from (Lawrence et al. 2006) and contained (in mM): 110 K-gluconate, 4 MgCl_2_, 0.1 EGTA, 4 Na_2_-ATP, 0.5 Na_2_-GTP, 10 Hepes, 10 Phosphocreatine (pH 7.3, 250 mOsm/L, 15 mV junction potential). Whole-cell voltage-clamp recording from mitral and tufted cells were made with an internal solution containing (in mM): 120 Cs-MeSO3, 20 tetraethylammonium-Cl, 5 4-aminopyridine, 2 MgCl_2_, 0.025 CaCl_2_, 1 EGTA, 4 Na-ATP, 0.5 Na-GTP, and 10 HEPES, (pH 7.3, 280 Osm/L, 10 mV junction potential). Atto 594 (10 µM, Sigma) was systematically added to the internal solution in order to visualize the cell morphology during the recording. Optical stimulation of the BF axons was done using a blue LED (490 nm, pE 100, CoolLED Ltd., Andover, UK) directed through the 40X objective of the microscope at 50-100% of its maximum power (5 mW at the objective output) and driven by the Axograph X acquisition software (Axograph Scientific). Olfactory nerves projecting inside a given glomerulus were electrically stimulated using a theta pipette filled with ACSF. The electrical stimulus (100 µs) was delivered using a Digitimer DS3 (Digitimer, Welwyn Garden City, UK). Recordings were low-passed filtered at 2-4 kHz and digitized at 20 kHz using the AxoGraph X software. In whole-cell voltage-clamp recordings, access resistance was not compensated. Voltages indicated in the paper were corrected for the junction potential.

### Cell selection

PG cells were selected based on the small size of their cell body and their position within the first rings of cells surrounding the glomerulus. Although they have a larger cell body, it cannot be excluded that TH-expressing cells or eTC have been erroneously included in the LCA recording dataset. However, these cells usually have a remarkable spontaneous activity patterns (eTC are rhythmically bursting, TH+ cells have a highly regular rhythmic discharge) and cells with this kind of activity constituted a minority of the dataset. eTC, superficial or middle tufted cells (collectively called s/mTC) and mitral cells were identified based on the localization of their soma and the presence or not of lateral dendrites in the external plexiform layer, as seen during whole-cell recording by visual inspection of the dye-filled cell morphology. In addition, some cells were filled with biocytin for post-hoc anatomical reconstruction. Thus, eTC were selected based on their large pear-shaped soma within the glomerular layer, a short and thick apical dendrite extensively ramifying into a single glomerulus and the lack of lateral dendrites. In addition, they often, but not always, spontaneously fired short burst of action potentials in the cell-attached configuration even in the presence of NBQX, d-AP5 and mecamylamine. Superficial tufted cells were found at the border between the glomerular layer and the external plexiform layer. Compared to eTC, their soma was located further from the glomerulus into which they projected and they had long lateral dendrites extending into the external plexiform layer. Middle tufted cells and mitral cells had large cell bodies located in the external plexiform layer and in the mitral cell layer, respectively, a thick apical dendrite projecting into a single glomerulus and long lateral dendrites in the external plexiform layer.

### Morphological reconstruction

Neurobiotin or biocytin (Vector Laboratories INC., Burlingame, CA) was added to the intracellular solution (1mg/ml). The patch pipette was slowly retracted after the recording to avoid damaging the cell body. The slice was then fixed in 4% paraformaldehyde overnight, washed 3 times in PBS and incubated in a permeabilizing solution containing Alexa Fluor 555 conjugated streptavidin (1 µg/ml; Thermo Fischer Scientific, Waltham, MA) overnight. After 3 rinses with PBS, sections were mounted in Vectashield Hardset with DAPI (Vector laboratories, Inc., Burlingame, CA). Labeled cells were imaged with a confocal microscope (Leica TCS SP5 II).

### Immunohistochemistry

ChAt mice expressing ChR2-eYFP in BF neurons were deeply anesthetized with Zolazepam tiletamine/Xylasine and transcardially perfused with PBS at room temperature followed by 4% paraformaldehyde (PFA, 4°C). Brains were removed, postfixed 3-6 hrs in 4% PFA at 4°C, rinsed in PBS and incubated in PBS until cut on a vibratome (VT 1000S, Leica). 50 µm-thick coronal sections were collected and stored in PBS. For ChAT staining, sections were incubated overnight at 4°C with a goat anti-choline acetyltransferase antibody (ChAT; 1:1000; Millipore Cat# AB144P, RRID:AB_2079751) in Tris-Triton buffer containing 2% donkey serum and 0.2% Triton X100. After three washes in Tris-Triton, sections were incubated 1 hour at room temperature with Alexa fluor 647 conjugated donkey anti-goat (1:500; ThermoFisher A-21447). After three washes, sections were mounted in Prolong Diamond Antifade Mountant (ThermoFisher) or Vectashield Hardset with DAPI. Images were taken using a Leica TCS SP5 II confocal microscope or a Zeiss Axio Imager M2 for mosaic images. Immunostained and EYFP-expressing cells were manually counted using the cell counter plug-in on Fiji software.

### Drugs

Chemicals used to prepare cutting, recording and internal solutions were acquired from MilliporeSigma, Carl Roth and Fisher Scientific. 6-nitro-7-sulfamoylbenzo[f]quinoxaline-2,3-dione (NBQX), D-2-Amino-5-phosphonopentanoic acid (D-AP5), 2-(3-carboxypropyl)-3-amino-6-(4 methoxyphenyl)pyridazinium bromide (gabazine), mecamylamine hydrochloride and atropine were purchased from Abcam Biochemicals. Scopolamine hydrobromide, Pirenzepine and XE-991 were purchased from Tocris Bioscience.

### Electrophysiological analysis

Action potential capacitive currents were automatically detected by the Axograph X software using an amplitude threshold. The timing of each spike was used to construct PSTH (peri stimulus time histograms) representing the total number of action potentials per time period (bin of 20 ms for 2 s-long episodes, of 200 ms for 15 s or longer episodes) across several consecutive sweeps (>30 for 2 s-long episodes, >10 for 15 s-long episodes). For cell classification, a post-stimulus spiking inhibition was a statistically significant (paired t-test or paired Wilcoxon signed-rank sum test) decrease in spike rate within a 100 ms or 200 ms-long time period immediately after the flash compared to the same period preceding the flash across at least 30 consecutive trials. The duration of spiking inhibition was calculated as the average duration between the flash and the first spike fired after the flash. A muscarinic excitation was a statistically significant increase in spike probability within a 1 s period starting 1 s after the flash compared to the 1s period immediately preceding the flash across at least 10 consecutive trials.

Photo-evoked IPSC amplitudes were measured as the peak of an average response computed from multiple sweeps. The decay of photo-evoked IPSCs was most often best fitted with a double exponential with time t=0 at the peak of the current. Time constant values indicated in the text are weighted decay time constants calculated using the following equation: τ_w_ = (τ_1_A_1_ + τ_2_A_2_)/(A_1_ + A_2_) where τ_1_ and τ_2_ are the fast and slow decay time constants and A_1_ and A_2_ are the equivalent amplitude weighting factors.

EPSCs and IPSCs were automatically detected by the Axograph X software using a sliding template function. To estimate the duration of an ON-evoked plurisynaptic excitatory response, PSTH representing the cumulative number of EPSCs per 20 ms bin across several consecutive sweeps were constructed. I then determined the time needed after stimulation for the EPSC frequency to come back to baseline frequency + 2 SD during at least five consecutive bins. Baseline frequency was calculated over the 25 bins (i.e. 500 ms) preceding the stimulation.

Data are presented as mean ± SD in the text and as mean ± SEM in graphs for display purpose. Data points from experiments were tested for normality using a Shapiro–Wilk test. Experiments with a normal distribution were tested for statistical significance with a paired Student’s t-test. Experiments with skewed distributions were tested for statistical significance using a paired Wilcoxon signed-rank sum test. For experiments comparing data points from different cells, statistical significance was determined using an unpaired t-test (normal distribution) or a Mann–Whitney test (non normal distribution).

## Aknowledgments

This work was supported by the Centre National de la Recherche Scientifique and the Université de Strasbourg. I thank Alvaro Sanz Diez, Julie Perraud and Andréa Grinner who contributed preliminary data. Ipek Yalcin and Matilde Cordero-Erausquin (Institut des Neurosciences Cellulaires et Intégratives, Strasbourg) for the kind gift of the dlx5/6-Cre mice and ChAT-Cre mice. Marion Najac, Philippe Isope, Antoine Valera, Karin Aubrey and Yo Otsu for their helpful comments on the manuscript. I also thank the staff of the animal facility (Chronobiotron, UMS 3415 CNRS and Strasbourg University) for technical assistance and members of the team Physiology of Neural Networks for their support throughout the project.

## Competing interests

The author declares that no competing interests exist.

## Supplementary figures

**Figure 3-figure supplement 1:**
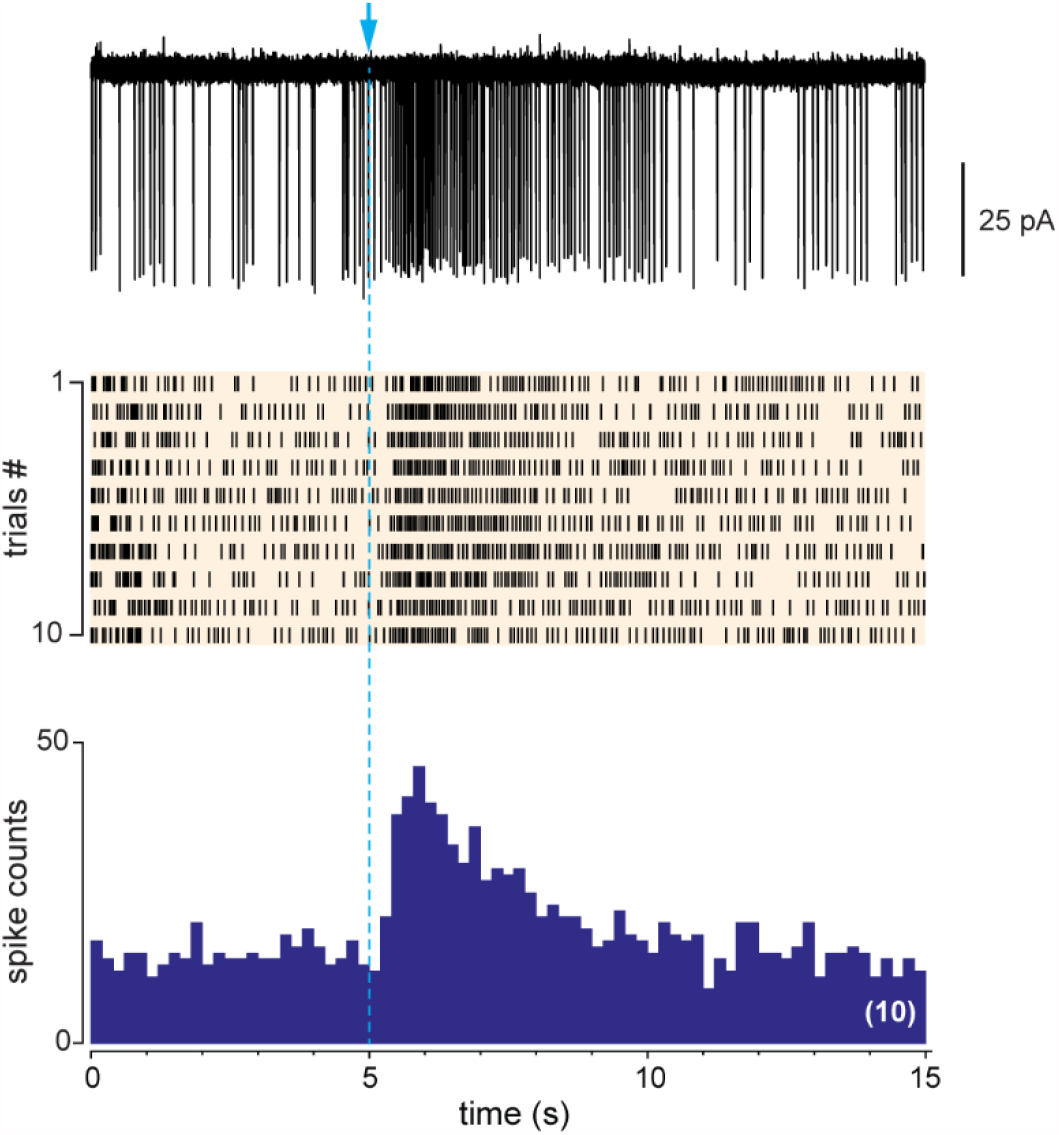
the unique example, in dlx5/6 mice, of a cell responding with a long-lasting excitation that was not accompanied by a transient inhibitory phase immediately after the flash

**Figure 7 – figure supplement 1:**
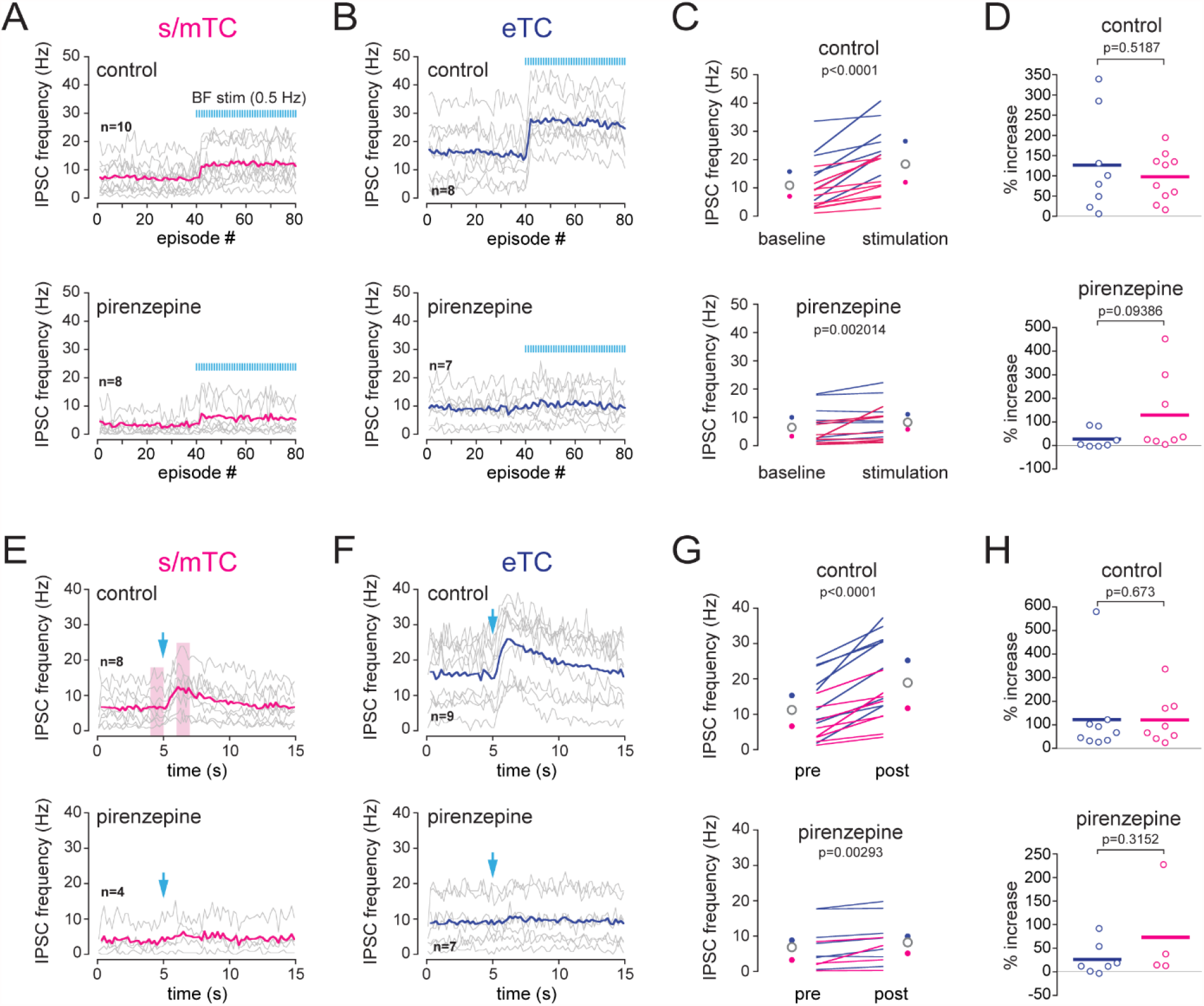
Comparison of BF-induced increase of inhibition in external tufted cells (eTC) and in superficial and middle tufted cells (s/mTC). **A-D**, Impact of a low frequency (0.5 Hz) photostimulation of the BF cholinergic axons on IPSC frequency in s/mTC and eTC in control condition (top) and in the presence of 2 µM pirenzepine (bottom). **A and B**, average IPSC frequency per episode (2 s each). Photostimulation of BF axons (1 flash/episode) starts at episode 41. Each line is a cell, colored lines are ensemble average. **C**, average IPSC frequency per cell when BF axons were not stimulated (baseline) and when BF cholinergic afferents were stimulated at 0.5 Hz (stimulation). Blue circles are the mean for eTC, magenta circles are the mean for s/mTC and opened circles are the ensemble means. **D**, percent BF-induced increase of ISPC frequency in eTC vs. in s/mTC. Horizontal lines are the average. **E-G**, Impact of a single photostimulation of the BF cholinergic axons on IPSC frequency in s/mTC and eTC in control condition (top) and in the presence of pirenzepine (bottom). The boxed areas in E show the pre- and post-stimulus time periods that were compared in G and H. Data points were all collected in slices from ChAT mice in the presence of NBQX, d-AP5 and mecamylamine (control condition).

**Figure 8 – figure supplement 1:**
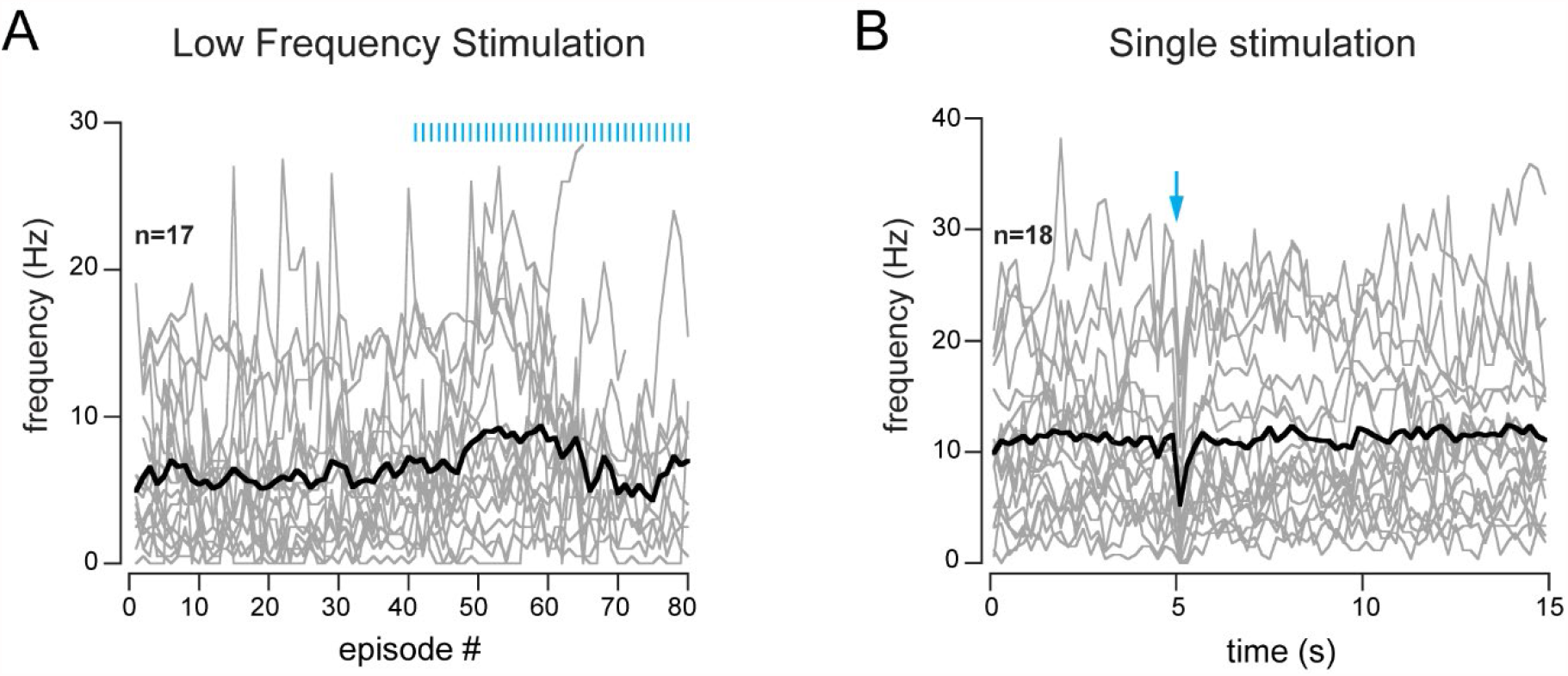
Absence of cholinergic excitation in PG cells showing a transient inhibition of spiking. **A**, low frequency stimulation of the BF axons (1 flash per 2 s-long episode, starting at episode 41). **B**, single photostimulation (blue arrow). Each grey line is a cell, the black line is the ensemble average.

**Figure 8 – figure supplement 2:**
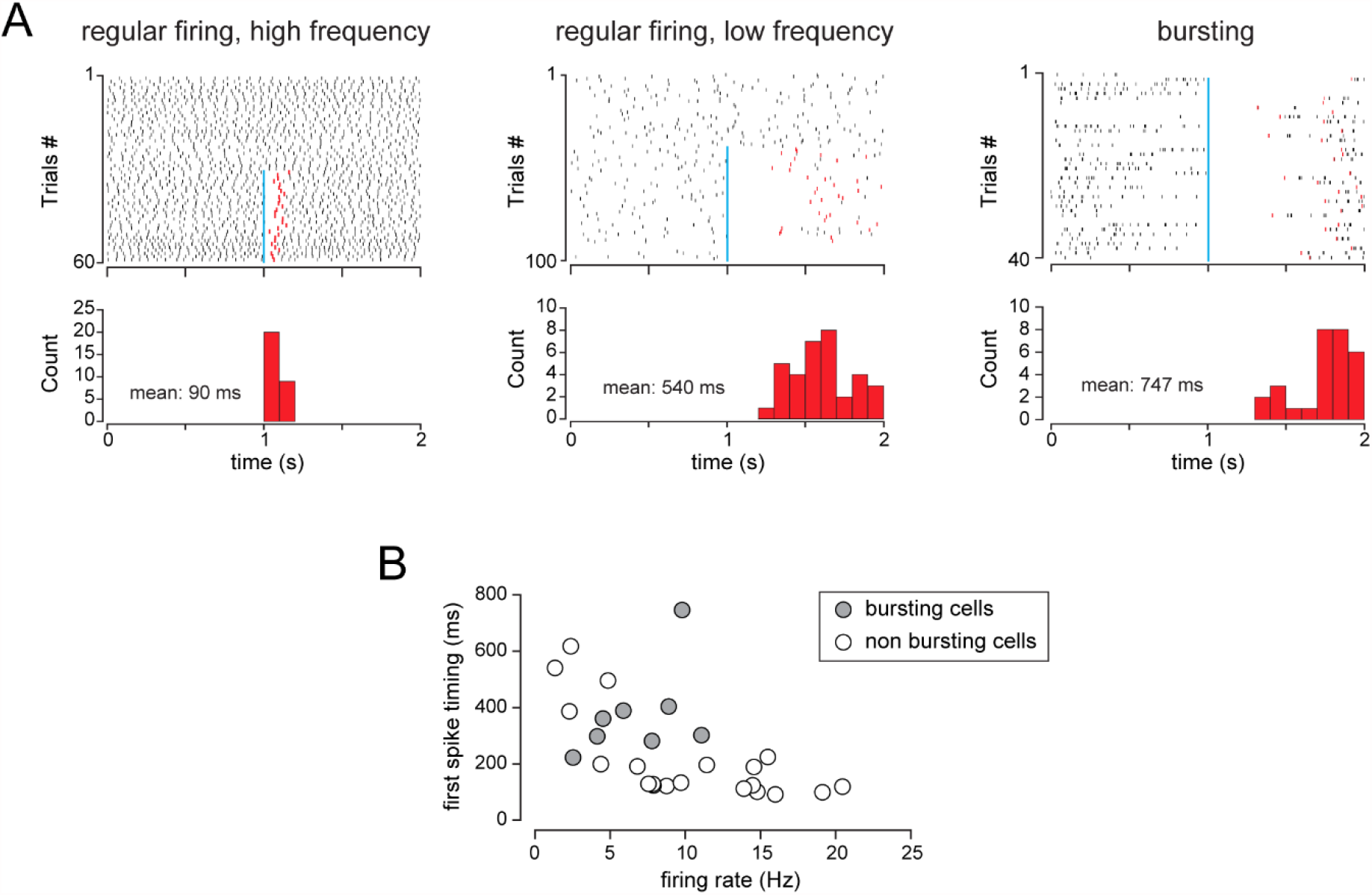
Caveats for the measurements of spiking inhibition duration. **A**, Raster plots from 3 example cells with various spiking activities that were inhibited by the BF input. The duration of the BF-induced inhibition was measured as the average time between the flash (blue bar) and the first spike after the flash (red tick). Histograms show the distribution of the first spike timing in each case and the calculated average duration. **B**, Average first spike timing in all the cells used for Fig.8A as a function of their baseline firing frequency.

## Notes

### Competing Interest Statement

The authors have declared no competing interest.

